# Inference of Adaptive Shifts for Multivariate Correlated Traits

**DOI:** 10.1101/146191

**Authors:** Paul Bastide, Cécile Ané, Stéphane Robin, Mahendra Mariadassou

## Abstract

To study the evolution of several quantitative traits, the classical phylogenetic comparative framework consists of a multivariate random process running along the branches of a phylogenetic tree. The Ornstein-Uhlenbeck (OU) process is sometimes preferred to the simple Brownian Motion (BM) as it models stabilizing selection toward an optimum. The optimum for each trait is likely to be changing over the long periods of time spanned by large modern phylogenies. Our goal is to automatically detect the position of these shifts on a phylogenetic tree, while accounting for correlations between traits, which might exist because of structural or evolutionary constraints. We show that, in the presence shifts, phylogenetic Principal Component Analysis (pPCA) fails to decorrelate traits efficiently, so that any method aiming at finding shift needs to deal with correlation simultaneously. We introduce here a simplification of the full multivariate OU model, named scalar OU (scOU), which allows for noncausal correlations and is still computationally tractable. We extend the equivalence between the OU and a BM on a re-scaled tree to our multivariate framework. We describe an Expectation Maximization algorithm that allows for a maximum likelihood estimation of the shift positions, associated with a new model selection criterion, accounting for the identifiability issues for the shift localization on the tree. The method, freely available as an R-package (`PhylogeneticEM`) is fast, and can deal with missing values. We demonstrate its efficiency and accuracy compared to another state-of-the-art method (ℓ**1ou**) on a wide range of simulated scenarios, and use this new framework to re-analyze recently gathered datasets on New World Monkeys and *Anolis* lizards.

## Motivation

A major goal of comparative and evolutionary biology is to decipher the past evolutionary mechanisms that shaped the present day diversity. Taking advantage of the increasing amount of molecular data made available by powerful sequencing techniques, sophisticated mathematical models have made it possible to infer reliable phylogenetic trees for ever growing groups of taxa (see e.g. Meredith et al. 2011; Jetz et al. 2012). Models of phenotypic evolution for such large families need to cope with the heterogeneity of observed traits across the species tree. One source of heterogeneity is the mechanism of “evolution by jumps” as hypothesized by Simpson (1944). It states that there exists an adaptive landscape shaping the evolution of functional traits, and that this landscape might shift, sometimes in a dramatic fashion, in response to environmental changes such as migration, or colonization of a new ecological niche. Such shifts, like the one observed in the brain shape and size of New World Monkeys in association with dietary and locomotion changes (Aristide et al. 2015, 2016), need to be explicitly accounted for in models of phenotypic evolution.

To detect such adaptive shifts, we must cope with two constraints: species do not evolve independently (Felsenstein 1985) and adaptive evolution is an intrinsically multivariate phenomenon. The first constraint arises from the shared evolutionary history of species, usually represented as a phylogenetic tree. It means that traits observed on closely related taxa are on average more similar than traits observed on distantly related species. The second constraint results from natural selection acting on many traits at once. Functional traits are indeed often interdependent, either because they are regulated by the same portions of the genetic architecture or because they are functionally constrained (e.g. limb bones lengths in Greater Antillean *Anolis* lizards Mahler et al. (2010)).

This work aims to develop a likelihood-based method to detect rapid adaptive events, referred to as shifts, using a time calibrated phylogenetic tree and potentially incomplete observations of a multivariate functional trait at the tips of that tree. The shifts can be used to cluster together species sharing a common adaptive history.

## State of the Art

Phylogenetic comparative methods (PCM) are the *de facto* tools for studying phenotypic evolution. Most of them can be summarized as stochastic processes on a tree. Specifically, given a rooted phylogeny, the traits evolve according to a stochastic process on each branch of the tree. At each speciation event, one independent copy with the same initial conditions is created for each daughter species. A common stochastic process in this setting is the Brownian Motion (BM, Felsenstein 1985). It is well suited to model the random drift of a quantitative, neutral and polygenic trait (see e.g. Felsenstein 2004, chap. 24). Unfortunately, the BM has no stationary distribution and cannot adequately model adaptation to a specific optimum (Hansen and Orzack 2005). The Ornstein-Uhlenbeck (OU) process is therefore preferred to the BM in the context of adaptive evolution (Hansen 1997; Hansen et al. 2008). Note that, as pointed out by Hansen et al. (2008) and Cooper et al. (2016), this model is distinct from the process theoretically derived by Lande (1976) for stabilizing selection toward an optimum on an adaptive landscape at a micro-evolutionary timescale, and is better seen as a heuristic for the macro-evolution of the “secondary optima” themselves in a Simpsonian interpretation of evolution (Hansen et al. 2008). Recently, Levy processes have also been used to capture Simpsonian evolution (Landis et al. 2013; Duchen et al. 2017).

Extensions to multivariate traits have been proposed for both BM (Felsenstein 1985) and OU processes (Bartoszek et al. 2012). Cybis et al. (2015) considered even more complex models, with a mix of both quantitative and discrete characters modeled with an underlying multivariate BM and a threshold model (Felsenstein 2005, 2012) for drawing discrete characters from the underlying continuous BM.

The work on adaptive shifts also enjoyed a growing interest in the last decade. In their seminal work, Butler and King (2004) considered a univariate trait with known shift locations on the tree and estimated shift amplitudes in the trait optimal value using a maximum-likelihood framework. Beaulieu et al. (2012) extended the work by estimating shift amplitudes not only in the optimal value but also in the evolutionary rate. The focus then moved to estimating the number and locations of shifts. Eastman et al. (2011, 2013) detected shifts, respectively, in the evolutionary rate or the trait expectations, for traits evolving as BM, in a Bayesian setting using reversible jump Markov Chain Monte Carlo (rjMCMC). Ingram and Mahler (2013); Uyeda and Harmon (2014); Bastide et al. (2016) detected shifts in the optimal value of a trait evolving as an OU. Uyeda and Harmon (2014) and Bastide et al. (2016) detect all shifts for a given number of shifts and use either rjMCMC or penalized likelihood to select the number of shifts. By contrast, Ingram and Mahler (2013) uses a stepwise procedure, based on AIC, to detect shifts sequentially, stopping when adding a shift does not improve the criteria anymore.

Extensions from univariate to multivariate shifts are more recent. It should be noted that all methods assume that shifts affect all traits simultaneously. Given known shift locations and a multivariate OU process, Bartoszek et al. (2012) was the first to develop a likelihood-based method (package `mvSLOUCH`) to estimate both matrices of multivariate evolutionary rates and selection strengths. Clavel et al. (2015) soon followed with `mvmorph`, a comprehensive package covering a wide range of multivariate processes. Detection of shifts in multivariate traits is more involved and both Ingram and Mahler (2013) and Khabbazian et al. (2016) make the simplifying assumption that all traits are independent, conditional on their shared shifts. Ingram and Mahler (2013) then proceed with the same stepwise procedure as in the univariate case whereas Khabbazian et al. (2016) uses a lasso-regression to detect the shifts and a phylogenetic BIC (pBIC) criterion to select the number of shifts.

## Scope of the Article

In this work, we present a new likelihood-based method to detect evolutionary shifts in multivariate OU models. We make the simplifying assumptions that all traits have the same selection strength but, unlike in Khabbazian et al. (2016) and Ingram and Mahler (2013), traits can be correlated. Our contribution is multifaceted. We show that the scalar assumption that we make (see Section Model) and the independence assumption share a similar feature in their structure that make the shift detection problem tractable. Building upon a formal analysis made in the univariate case (Bastide et al. 2016), we show that the problem suffers from identifiability issues as two or more distinct shift configurations may be indistinguishable. We propose a latent variable model combined with an OU to BM reparametrization trick to estimate the unknown number of shifts and their locations. Our method is fast and can handle missing data. It also proved accurate in a large scale simulation study and was able to find back known shift locations in re-analysis of public datasets. Finally, we show that the standard practice of decorrelating traits using phylogenetic principal component analysis (pPCA) before using a method designed for independent traits can be misleading in the presence of shifts.

The article is organized as followed. We present the model and inference procedure in Section Model, the theoretical bias of pPCA in the presence of shifts in Section pPCA and Shifts, the simulation study in Section Simulations Studies, the re-analysis of the New World Monkeys and Greater Antillean *Anolis* lizards datasets in Section Examples and discuss the results and limitations of our method in Section Discussion.

## Model

### Trait Evolution on a Tree

*Tree*.— We consider a fixed and time-calibrated phylogenetic tree linking the present-day species studied. The tree is assumed ultrametric with height *h*, but with possible polytomies. We denote by *n* the number of tips and by *m* the number of internal nodes, such that *N* = *n* + *m* is the total number of nodes. For a fully bifurcating tree, *m* = *n* — 1, and *N* = 2*n* — 1.

*Traits*.— We note **Y** the matrix of size *n* × *p* of measured traits at the tips of the tree. For each tip *i*, the row-vector **Y**^*i*^ represents the *p* measured traits at tip *i*. Some of the data might be missing, as discussed later (see Section Statistical Inference).

*Brownian Motion* (*BM*).— The multivariate BM has *p* + *p*(*p* + 1)/2 parameters: *p* for the ancestral mean value vector *μ*, and *p*(*p* + 1)/2 for the drift rate (in the genetic sense) matrix **R**. The variance of a given trait grows linearly in time, and the covariance between two traits *k* and *l* at nodes *i* and *j* is given by *t*_*ij*_*R*_*kl*_, where *t*_*ij*_ is the time elapsed between the root and the most recent common ancestor (MRCA) of *i* and *j* (see e.g. Felsenstein 2004, chap. 24). Using the vectorized version of matrix **Y** (where vec(**Y**) is the vector obtained by “stacking” all the columns of **Y**), we get: var [vec(**Y**)] = **R** ⊗ **C**, where ⊗ is the Kronecker product, and **C** = [*t*_*ij*_]_1*≤i,j*≤*n*_.

*Ornstein-Uhlenbeck* (*OU*).— The Ornstein-Uhlenbeck process has *p*^2^ extra parameters in the form of a selection strength matrix **A**. The traits evolve according to the stochastic differential equation *d***X**_*t*_ = **Α**(*β* — **X**_*t*_)*dt* + **R***d***W**_*t*_, where **W**_*t*_ stands for the standard *p*-variate Brownian motion. The first part represents the attraction to a “primary optimum” *β*, with a dynamic controlled by **A**. This matrix is not necessarily symmetric in general, but it must have positive eigenvalues for the traits to indeed be attracted to their optima. This assumption also ensures the existence of a stationary state, with mean *β* and variance **Γ** (see Bartoszek et al. 2012; Clavel et al. 2015, for further details and general expression of **Γ**).

*Shifts*.— We assume that some environmental changes affected the traits evolution in the past. In the BM model, we take those changes into account by allowing the process to be discontinuous, with shifts occurring in its mean value vector (as e.g. Eastman et al. 2013). This is reasonable if the adaptive response to a change in the environment is fast enough compared to the evolutionary time scale. For the OU, we assume that environmental changes result in a shift in the primary optimum *β* (as e.g. Butler and King 2004). The process is hence continuous, and goes to a new optimum, with a dynamic controlled by **A**. In both cases, we make the standard assumptions that all traits shift at the same time (but with different magnitudes), that each shift occurs at the beginning of its branch, and that all other parameters (**A, R**) of the process remain unchanged. We further assume that each jump induces a specific optimum, which implies that there is no homoplasy for the optimum, that is, no convergent evolution.

### Simplifying Assumptions

*Trait Independence Assumption*.— The general OU as described above is computationally hard to fit (Clavel et al. 2015), even when the shifts are fixed *a priori*. For automatic detection to be tractable in practice, several assumptions can be made. The two methods that (to our knowledge) tackle this problem in the multivariate setting assume that all the traits are independent, i.e. that matrices **A** and **R** are *diagonal* (Ingram and Mahler 2013; Khabbazian et al. 2016). This is often justified by assuming that *a priori* preprocessing with phylogenetic Principal Component Analysis (pPCA, Revell 2009) leads to independent traits. However, pPCA assumes a no-shift BM evolution of the traits, and it can introduce a bias in the downstream analysis conducted on the scores, as shown by Uyeda et al. (2015). The choice of the number of PC axes to keep is also crucial, and can qualitatively change the results obtained, leading to the detection of artificial shifts near the root when not enough PC axes are kept for the analysis, as observed by Khabbazian et al. (2016). Finally, we show theoretically (Section pPCA and Shifts) and numerically (Section Simulations Studies, last paragraph) that pPCA fails to decorrelate the data in the presence of shifts and may even hamper shift detection accuracy.

*Scalar OU* (*scOU*).— We offer here an alternative to the independence assumption. Computations are greatly simplified when matrices **A** and **R** commute. This happens when both of these matrices are diagonal for example, or when **R** is unconstrained and **A** is *scalar*, i.e. of the form **A** = *α***I**_*p*_, where **I**_*p*_ is the identity matrix. We call a process satisfying the latter assumptions a *scalar OU* (scOU), as it behaves essentially as a univariate OU. In particular, its stationary variance is simply given by **Γ** = **R**/(2*α*) (analogous to the formula *γ*^2^ = *σ*^2^/(2*α*) in the univariate case, see e.g. Hansen 1997).

We define the scOU model as follows: at the root *ρ*, the traits are either drawn from the stationary normal distribution with mean *μ* and variance **Γ** (**X**^*ρ*^ ~ 𝕍(*μ*, **Γ**)), or fixed and equal to *μ*. The initial optimum vector is *β*_0_ and the conditional distribution of trait **X**^*i*^ at node *i* given trait **X**^pa(*i*)^ at its parent node pa(*i*) is

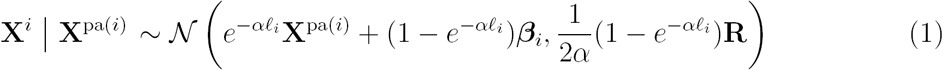

where *β*_*i*_ = *β*_Ρa(*i*)_ + **Δ**^*i*^ is the optimal value of the process on the branch with length ℓ_*i*_ going from pa(*i*) to *i* and **Δ** is the *N* × *p* matrix of shifts on the branches of the tree: for any node *i* and any trait *l*, **Δ**_*il*_ is 0 if there are no shift on the branch going from pa(*i*) to *i*, and the value of the shift on trait *l* otherwise. At the root, we define *β*_*ρ*_ = *β*_0_ and, for each trait *l*: **Δ**_*ρl*_ = *e*^−*αh*^*μ*_*l*_ + (1 — *e*^−*ah*^)*β*_0*l*_, where *h* is the age of the root (or tree height).

The scOU model can also be expressed under a linear form. Let **U** be the *N × N* matrix where *U*_*ij*_ is 1 if node *j* is an ancestor of node *i* and 0 otherwise. Let **T** be the *n × N* matrix made of the *n* rows of **U** corresponding to tip taxa. For a given *α*, we further define the diagonal *N* matrix **W**(*α*) with diagonal term *W*_*ii*_(*α*) = 1 — e^−*αα*pa(*i*)^ for any non-root node *i*, where *α*_pa(*i*)_ is the age of node pa(*i*), and *W*_*ρρ*_(*α*) = 1 for the root node _*ρ*_. Then the joint distribution of the observed traits **Y** is normal

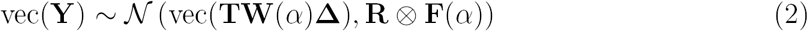

where **F**(*α*) is the symmetric scaled correlation matrix between the *n* tips, with entries 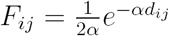if the root is drawn from the stationary distribution, and 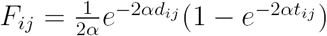if the root is fixed, where *d*_*ij*_ is the tree distance between nodes *i* and *j*. In the next section, this will allow us to rewrite scOU as a BM on a tree with rescaled branch lengths. This observation is at the core of our statistical inference strategy.

The scOU process allows us to handle the correlations that might exist between traits, and spares us from doing a preliminary pPCA. This however comes at the cost of assuming that all the traits evolve at the same rate toward their respective optima, with the same selection strength *α*. See the Discussion for further analysis of these assumptions.

### Identifiability Issues

*Root State*.— It can be easily checked that the parameters ***μ*** and ***β***_0_ at the root are not jointly identifiable from observations at the tips of an ultrametric tree, only the combination **λ** = *e*^−*αh*^***μ*** + (1 — *e*^−*ah*^)***β***_0_ is. See Ho and Ané (2014) for a derivation in the univariate case. Note that **λ** corresponds to the first row of the shift matrix **Δ**. As we cannot decide from the data, we assume by default ***β***_0_ = ***μ*** = **λ**.

*Shift Position*.— The location of the shifts may not always be uniquely determined, as several sets of locations (and magnitudes) may yield the same joint marginal distribution of the traits at the tips. These identifiability issues have been carefully studied in Bastide et al. (2016) for the univariate case. Because we assume that all traits shift at the same time, the sets of equivalent shift locations are the same in the multivariate case as in the univariate case; only the number of parameters involved is different. So, the problem of counting the total number of parsimonious, non-equivalent shift allocations remains the same, as well as the problem of listing the allocations that are equivalent to a given one.

As a consequence, all the combinatorial results and algorithms used in Bastide et al. (2016) are still valid here; only the model selection criterion needs be adapted (see Section Statistical Inference).

### Re-scaling of the Tree

*Equivalency scOU / rBM*.— As recalled above, the inference of OU models raises specific issues, mostly because some maximum likelihood estimates do not have a closed form expression. Many of these issues can be circumvented using the equivalence between the univariate BM and OU models described in Blomberg et al. (2003); Ho and Ané (2013); Pennell et al. (2015), for ultrametric trees, when α is known. Thanks to the scalar assumption, this equivalence extends to the multivariate case. Indeed, the marginal distribution of the traits at the observed tips **Y** given in (2) is the same as the one arising from a BM model on a re-scaled tree defined by:

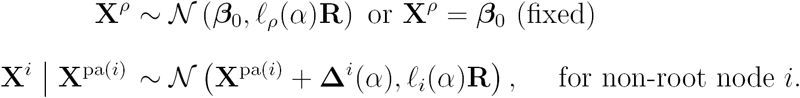

where 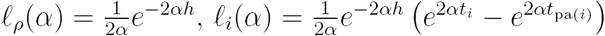, and 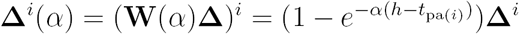. Note that, when the root is taken random, everything happens as if we added a fictive branch above the root with length *ℓ*_*ρ*_(*α*). The length of this branch increases when α goes to zero.

We emphasize that only the distribution of the observed traits **Y** is preserved and not the distribution of the complete dataset **X**. As a consequence, ancestral traits at internal nodes cannot be directly inferred using this representation. Still, the equivalence recasts inference of **R** and **W**(*α*)**∆** in the scOU model into inference of the same parameters in a much simpler BM model, albeit on a tree with rescaled branch lengths *ℓ*_*i*_(*α*). Note that the rescaling depends on α, which needs to be inferred separately. See the discussion (Section Interpretation Issues) for further analysis of this re-scaling.

### Statistical Inference

*Incomplete Data Model*.— We now discuss how to infer the set of parameters ***θ*** = (**Δ, R**). We adopt a maximum likelihood strategy, which consists in maximizing the log-likelihood of the observed tip data log *p***_*θ*_**(**Y**) with respect to ***θ*** to get the estimate 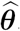. The maximum likelihood estimate 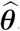is difficult to derive directly as the computation of log *p***_*θ*_**(**Y**) requires to integrate over the unobserved values of the traits at the internal nodes. We denote by **Z** the unobserved matrix of size *m × p* of these ancestral traits at internal nodes of the tree: for each internal node *j*, **Z**^*j*^ is the row-vector of the *p* ancestral traits at node *j*. Following Bastide et al. (2016), we use the expectation-maximization (EM) algorithm (Dempster et al. 1977) that relies on an incomplete data representation of the model and takes advantage of the decomposition of log *p***_*θ*_**(**Y**) as 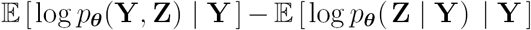.

*EM*.— The *M* step of the EM algorithm consists in maximizing 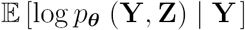with respect to ***θ***. For a given value of *α*, thanks to the rescaling described in Section Model, the formulas to update **Δ** and **R** are explicit (see Appendix EM Inference). The optimization of *α* is achieved over a grid of values, at each point of which a complete EM algorithm is run.

At the M step, we need the mean and variance of the unobserved traits **Z**^*j*^ at each internal node *j* conditional on the observed traits **Y** at the tips. The E step is dedicated to the computation of these values, which can be achieved via an upward-downward recursion (Felsenstein 2004). The upward path goes from the leaves to the root, computing the conditional means and variances at each internal node given the values of its offspring in a recursive way. The downward recursion then goes from the root to the leaves, updating the values at each internal node to condition on the full taxon set. Thanks to the joint normality of the tip and internal node data, all update formulas have closed form matrix expressions, even when there are some missing values (see Appendix EM Inference).

*Initialization*.— The EM algorithm is known to be very sensitive to the initialization. Following Bastide et al. (2016), we take advantage of the linear formulation (2) to initialize the shifts position using a lasso penalization (Tibshirani 1996). This initialization method is similar to the procedure used in *ℓ***1ou** (Khabbazian et al. 2016). See Appendix EM Inference for more details.

*Missing Data*.— EM was originally designed to handle missing data. As a consequence, the algorithm described above also applies when some traits are unobserved for some taxa. Indeed, the conditional distribution of the missing traits given the observed ones can be derived in the same way as in the E step. However, missing data break down the factorized structure of the dataset and some computational tricks are needed to handle the missing data efficiently (see Appendix EM Inference).

*Model Selection*.— For each value of the number of shifts *K*, the EM algorithm described above provides us with the maximum likelihood estimate 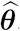. *K* needs to be estimated to complete the inference procedure. We do so using a penalized likelihood approach. The model selection criterion relies on a reformulation of the model in terms of multivariate linear regression, where we remove the phylogenetic correlation, like independent contrasts and PGLS do. We can re-write (2), for a given *α*, as

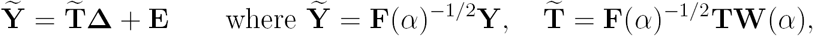

where **E** is a *n × p* matrix with independent and identically distributed rows, each row being a (transposed) centered Gaussian vector with variance **R**. In the univariate case (Bastide et al. 2016), this representation allowed us to cast the problem in the setting considered by Baraud et al. (2009), and hence to derive a penalty on the log-likelihood, or, equivalently, on the least squares. Taking advantage of the well known fact that the maximum likelihood estimators of the coefficients are also the least square ones, and do not depend on the variance matrix **R** (see, e.g. Mardia et al. 1979, Section 6), we propose to estimate *K* using the penalized least squares:

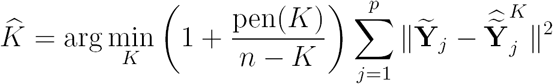

where 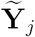is the column of 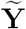 for the *j*-th trait, and 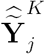the predicted means for trait *j* from the best model with *K* shifts. Using the EM results, this can be written as:

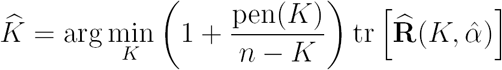

where 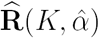is the ML estimate of the variance parameter obtained by the EM for a fixed number *K* of shifts. The penalty is the same as in the univariate case:

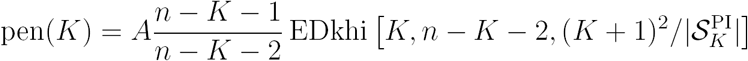

where EDkhi is the function from Definition 3 from Baraud et al. (2009) and 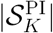is the number of parsimonious identifiable sets of locations for *K* shifts, as defined in Bastide et al. (2016). It hence might depends on the topology of the tree, for a tree with polytomies. For a fully resolved tree, 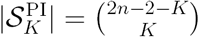. A is a normalizing constant, that must be greater than 1. In Baraud et al. (2009), the authors showed that it had little influence in the univariate case, and advised for a value around *A* = 1.1. We took this value as a default.

The criterion is directly inspired from the univariate case and inherits its theoretical properties in the special case **R** = *σ***^2^Ι**_*p*_. In general however, the criterion should be seen as a heuristic, although with good empirical properties (see Section Simulations Studies).

### Implementation

We implemented the method presented above in the **PhylogeneticEM R** package (R Core Team 2017), available on the Comprehensive R Archive Network (CRAN). A thorough documentation of its functions, along with a brief tutorial, is available from the **GitHub** repository of the project **(pbastide.github.io/PhylogeneticEM)**. Thanks to a comprehensive suite of unitary tests, that cover approximately 79% of the code **(codecov.io/gh/pbastide/PhylogeneticEM),** and that are run automatically on an independent Ubuntu server using the continuous integration tool **Travis CI (travis-ci.org),** the package was made as robust as possible. The computationally intensive parts of the analysis, such that the upward-downward algorithm of the M step, have been coded in **C++** to improve performance (see Section Simulations Studies for a study of the computation times needed to solve problems of typical size). Because the inference on each *α* value on the grid used is independent, they can be easily be done in parallel, and a built in option allows the user to choose the number of cores to be allocated to the computations.

## pPCA and Shifts

Shift detection in multivariate settings is usually done by first decorrelating traits with pPCA before feeding phylogenetic PCs to detection procedures that assume independent traits. We show hereafter that even in the simple BM setting, phylogenetic PC may still be correlated in the presence of shifts. The problem is only exacerbated in the OU setting.

### pPCA is biased in the presence of shifts

Assume that the traits evolve as a shifted BM process on the tree, so that 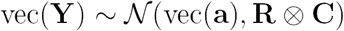, with **a** being the *n × p* matrix of trait means at the tips. Decomposing **R** as **R** = **VD^2^V^*T*^**, pPCA relies on the fact that the columns of the matrix **YV** are independent. Therefore, its efficiency relies on an accurate estimation of **R**.

The estimate of **R** used in pPCA is 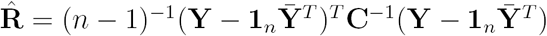, where 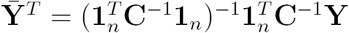, which is known as the estimated phylogenetic mean vector (Revell 2009). Decomposing the estimate of **R** as 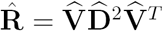, pPCA then computes the scores as 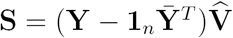.

In the absence of shift, all species have the same mean vector ***μ*** so **a** = 1_*n*_***μ*^*T*^** and 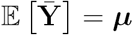. In the presence of shifts, species do not all share the same mean vector so the uniform centering is not valid anymore. As a consequence, the estimate of **R** is biased (see appendix PCA: Mathematical Derivations):

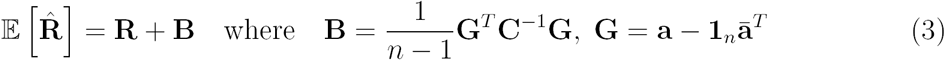

The extra term **B** is analogous to the between-group variance in the context of linear discriminant analysis and cancels out in the absence of shifts (note that **R** is analogous to the within-group variance, see Mardia et al. 1979). Because 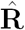 is biased, the columns of the score matrix **S** resulting from pPCA are still correlated. We illustrate this phenomenon below using toy examples.

### Illustration: a simple example

To illustrate the impact of shifts on the decorrelation performed by (p)PCA, we used the simple tree presented in Figure 1a and considered three scenarios. In all scenarios, we simulated two highly correlated traits under a BM starting from (0, 0) at the root and with covariance matrix 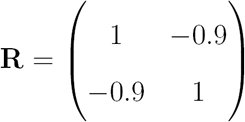. The tree has two clearly marked clades, designed to highlight the differences between pPCA and PCA. **R** is identical in all scenarios; any preprocessing aiming at decorrelating the traits should retrieve the eigenvectors of **R** as PCs. In the first scenario, there are no trait shifts on the tree, corresponding to the pPCA assumptions, and pPCA is indeed quite efficient in finding the PCs (see Fig. 1b, left panel). In the second scenario, we added a shift on a long branch. This shift induces a species structure in the trait space that misleads standard PCA. The same structure can however be achieved by a large increment of the BM on that branch and large increments are likely on long branches. pPCA therefore copes with the shift quite well and is able to recover accurate PCs. More quantitatively, the bias induced by the shift on 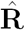 is quite small, 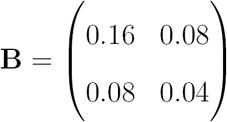, around one tenth of the values of **R**. In the third scenario, we put a shift on a small branch. The structure induced by the shift “breaks down” the upper clade and is unlikely to arise from the increment of a BM on that branch. It is therefore antagonistic to pPCA and results in a large bias for 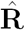: the extra term **B** is equal to 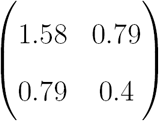and comparable to **R**. In that scenario, both PCA and pPCA find axes that are far away from the eigenvectors of **R** (Figure 1b, right panel). The first eigenvector of **R** captures the evolutionary drift correlation between traits, whereas the PCs of both PCA and pPCA capture a mix of evolutionary drift correlation and correlation resulting from shifts along the tree.

**Figure 1:**
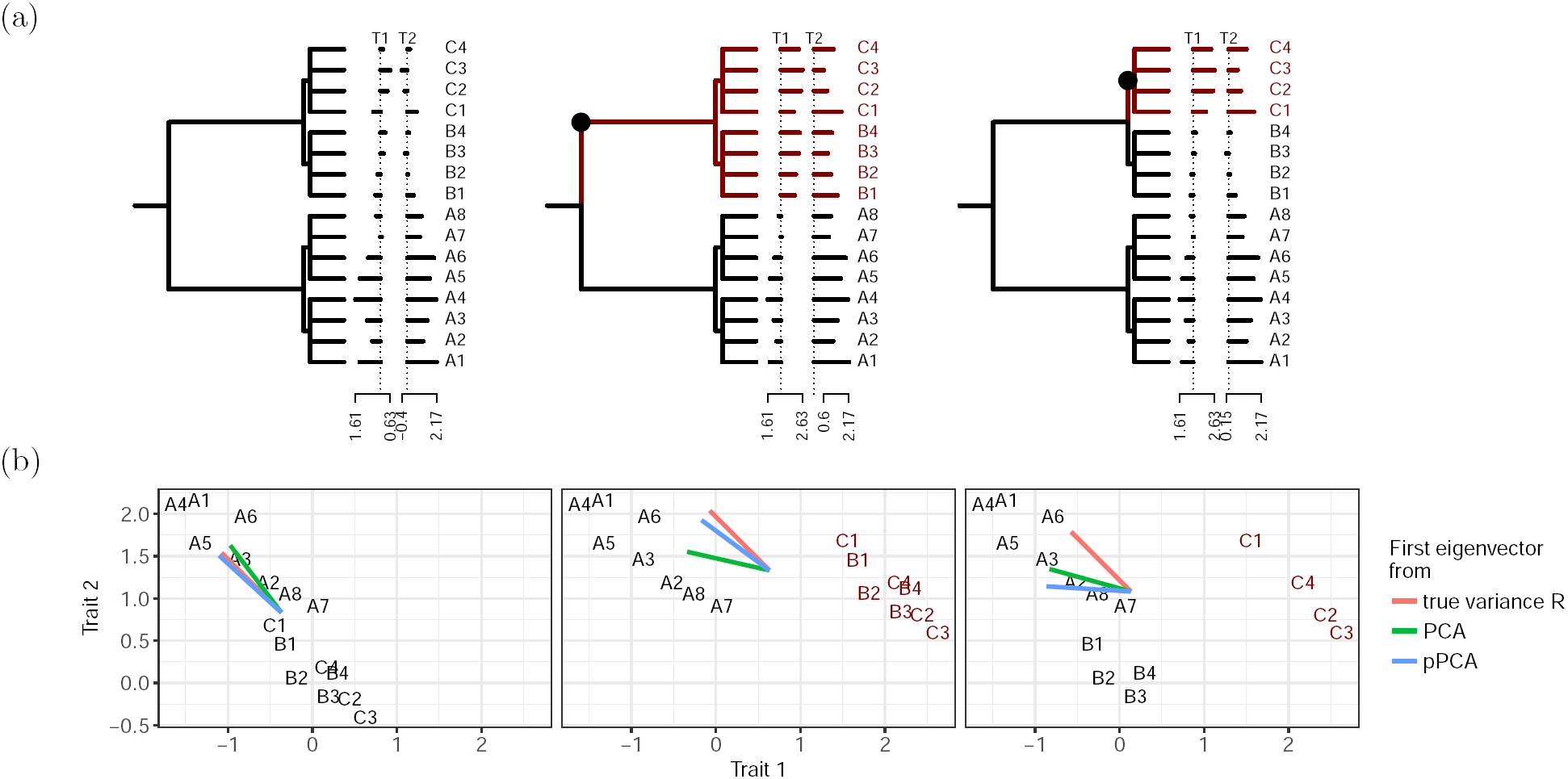
Bivariate traits simulated as a BM under three scenarios: no shift (left), shift on a long branch (middle) and shift on a short branch (right). Species affected by the shift are in dark red. Top: Phylogenetic tree, shift position and simulated trait values. Bottom: Scatterplot of species in the trait space and corresponding first eigenvector computed from the true covariance **R** (red) or found by PCA (green) and pPCA (blue).

## Simulations Studies

### Experimental Design

*General Setting*.— We studied the performance or our method using a “star-like” experimental design, as opposed to a full-factorial design. We first considered a base scenario, corresponding to a base parameter set, and then varied each parameter in turn to assess its impact as in Khabbazian et al. (2016). The base scenario was chosen to be only moderately difficult, so that our method would find shifts most but not all of the time.

For the base scenario, we generated one 160-taxon tree according to a pure birth process, using the **R** package **TreeSim** (Stadler 2011), with unit height and birth rate λ = 0.1. We then generated 4 traits on the phylogeny according to the scOU model, with a rather low selection strength *α*_*b*_ =1 (*t*_1/2_ = 69% of the tree height), and with a root taken with a stationary variance of 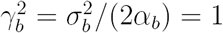. Diagonal entries of the rate matrix **R** are 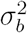and off-diagonal entries were set to 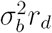with a base correlation of *r*_*d*_ = 0.4 (correlated traits) when testing the effect of shift number and amplitude, or *r*_*d*_ = 0 (independant traits) otherwise.

Finally, we added three shifts on this phylogeny, with fixed positions (see Figure 2). Shift amplitudes were calibrated so that the means at the tips differ by about 1 standard deviation, which constitute a reasonable shift signal (Khabbazian et al. 2016). Each configuration was replicated 100 times. We then used both our **PhylogeneticEM** and *ℓ***1ou** package (Khabbazian et al. 2016) to study the simulated data. We excluded **SURFACE** (Ingram and Mahler 2013) from the comparison at is (i) quite slow, (ii) assumes the same evolutionary model as *ℓ***1ou** and (iii) was found to achieve worse accuracy than *ℓ***1ou** (Khabbazian et al. 2016). We used default setting for both methods. For **PhylogeneticEM** this implies an inference on an automatically chosen grid with 10 *α* values, on a log scale, and a maximum number of shifts of 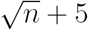(See Bastide et al. 2016 and Appendix EM Inference for a justification of these default parameters).

**Figure 2:**
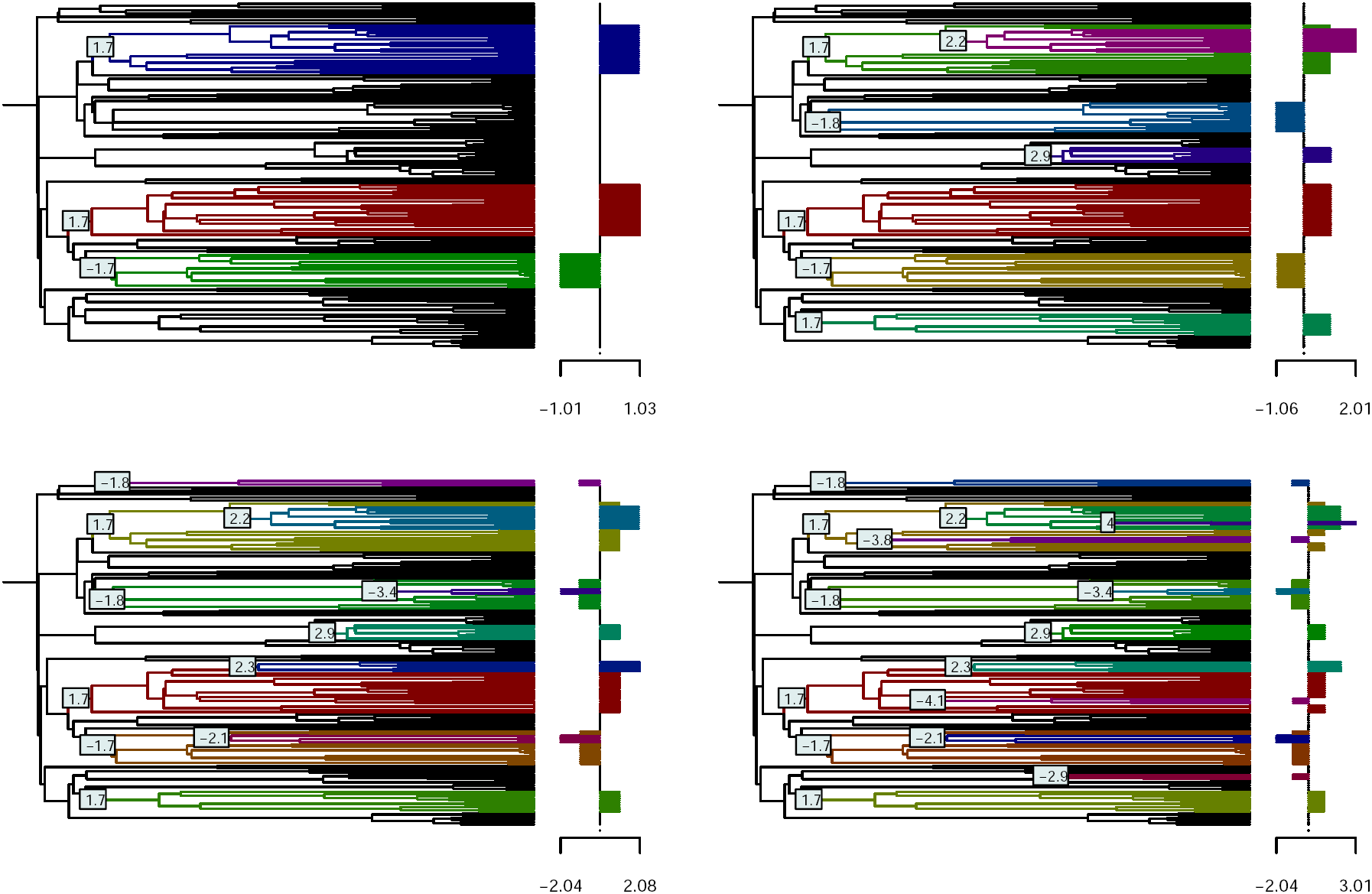
Shifts locations and magnitudes used in the base scenario. Mean trait values are identical for the 4 traits, up to a multiplicative ±1 factor and shown at the tips. Colors correspond to the different regimes. The bar plots on the right represent the expected traits values under the base model.

*Number and Amplitude of Shifts*.— We explored the effect of shifts by varying both their number and amplitude. We considered successively 0, 3, 7, 11, 15 shifts on the topology, with positions and values fixed as in Figure 2. Shifts values were chosen to form well separated tip groups; adjacent (in the tree) group means differ by about 1 standard deviation γ_*b*_. To mimic adaptive events having different consequences on different traits, all shifts on a trait were then randomly multiplied by —1 or +1. Finally and to assess the effect of shift amplitude, we rescaled all shifts by a common factor taking values in [0.5, 3]. Low scaling values correspond to smaller, harder to detect, shifts and high values to larger and easier to detect shifts.

*Selection Strength*.— When exploring parameters not related to the shifts, we considered a base number of 3 shifts and a base scaling factor of 1.25, empirically found to correspond to a moderately difficult scenario. We also assumed independent traits with the same variance and selection strength (i.e. scalar **A** and **R**, see *model A* in appendix Kullback-Leibler Divergences). We first varied *α* from 1 to 3 (i.e. *t*_1/2_ varied between 35% and 23% of the tree height). The variance *σ*^2^ varied with *α* to ensure that the stationary variance 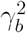remained fixed at 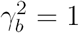.

*Model Mis-specification*.— The two current frameworks (*ℓ***1ou** and scOU) for multivariate shift detection assume independents traits (diagonal **A** and **R**) or correlated traits with equal selection strengths (scalar **A** and arbitrary **R**). To assess robustness to model mis-specification, we simulated data under four classes of models, referred to as A, B, C, D. Model A is correctly specified for both scOU and *ℓ***1ou** whereas B, C, D correspond respectively to mis-specifications for *ℓ***1ou,** scOU and both. We used the Kullback-Leibler divergence between models A and B (resp. C, D) to choose parameters that attain comparable “levels” of mis-specification (see appendix Kullback-Leibler Divergences for details).

- *Model A* assumes scalar **A** and **R** (independent traits, same selection strength and variance) and meets the assumptions of both scOU and *ℓ***1ou**.
- *Model B* assumes scalar **A** and arbitrary **R** (correlated traits, same selection strength) and corresponds to the scOU model. The level of correlation is controlled by setting all off-diagonal terms to 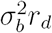in **R**. Following Khabbazian et al. (2016), *r*_*d*_ varies from 0.2 to 0.8, leading to Kullback divergences of up to 288.36 units.
- *Model C* assumes diagonal, but not scalar, **A**, and diagonal **R** (independent traits, different selection strengths), which matches the assumptions of *ℓ***1ou** only. We considered **A** = *α* Diag(s^−1.5^, s^−0.5^, s^0.5^, s^1.5^) with *s* varying from 2 to 8. We accordingly set 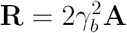to ensure that all traits have stationary variance 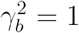. This led to Kullback divergences of up to 286.78 units.
- *Model D* assumes non-diagonal **A** and diagonal **R** (uncorrelated drift, but correlated traits selection) and violates both models. Following Khabbazian et al. (2016), all off-diagonal elements of **A** were set to *α*_*b*_*r*_*s*_ varying from 0.2 to 0.8. In this case, the stationary variance is not diagonal but has diagonal entries equal to 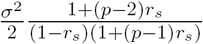. We thus rescaled *σ*^2^ appropriately to ensure that each trait has marginal stationary variance 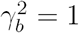as previously. This led to Kullback divergences of up to 112.98 units.

We expected *ℓ***1ou** to outperform scOU in model C and vice versa in model B. To be fair to both methods, we selected parameter ranges leading to similar Kullback divergences, to achieve similar levels of mis-specifications. However, both deviations produce datasets with groups that are also theoretically easier to discriminate compared to model A (see Figure 3). Indeed, we can quantify the difficulty of a dataset in terms of group separation by the Mahalanobis distance between the observed data and their expected mean, (phylogenetically) estimated in the absence of shifts:

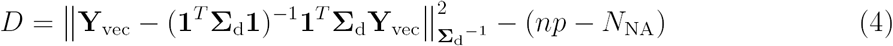

where **Y_vec_** is the vector of observed data at the tips (omitting missing values), **Σ_d_** is the true variance of **Y_vec_** and *N*_NA_ is the number of missing values. In the absence of shifts 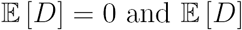increases when groups are well separated.

**Figure 3:**
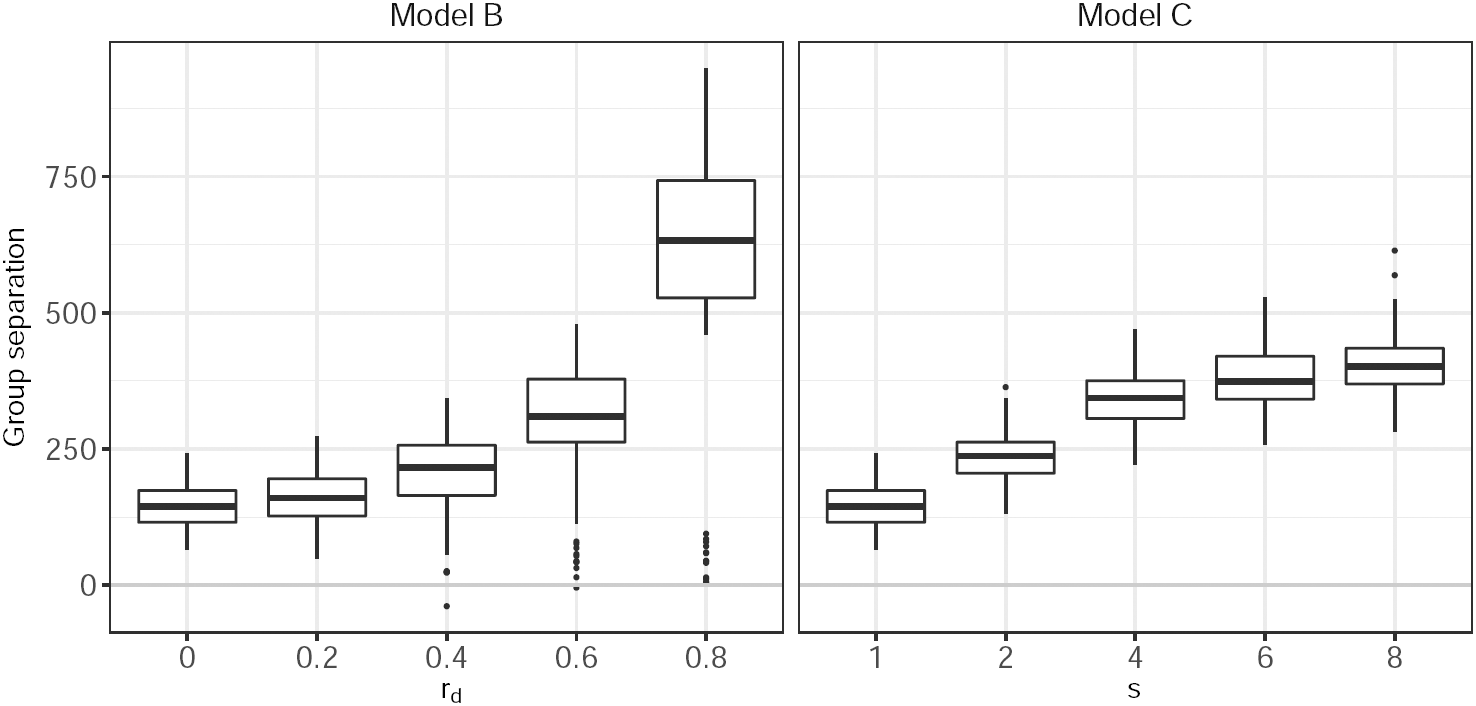
Impact of trait correlation *r*_*d*_ (left) and unequal selection strengths *s* (right) on group separation, as defined in Eq. (4). Unequal selection strengths (*s* > 1) and trait correlations (*r*_*d*_ > 0) both increase group separation and make it easier to detect shifts.

*Number of Observations*.— We varied the number of observations by (i) varying the number of taxa and (ii) adding missing values. To change the number of taxa, we generated 6 extra trees with the same parameters as before but with 32 to 256 taxa. The three shifts were fixed as in Figure 4. To test the ability of our method to handle missing data, we removed observations at random in our base scenario, taking care to keep at least one observed trait per species, so as not to change the number of taxa. The fraction of missing data varied from 5% to 50%.

**Figure 4:**
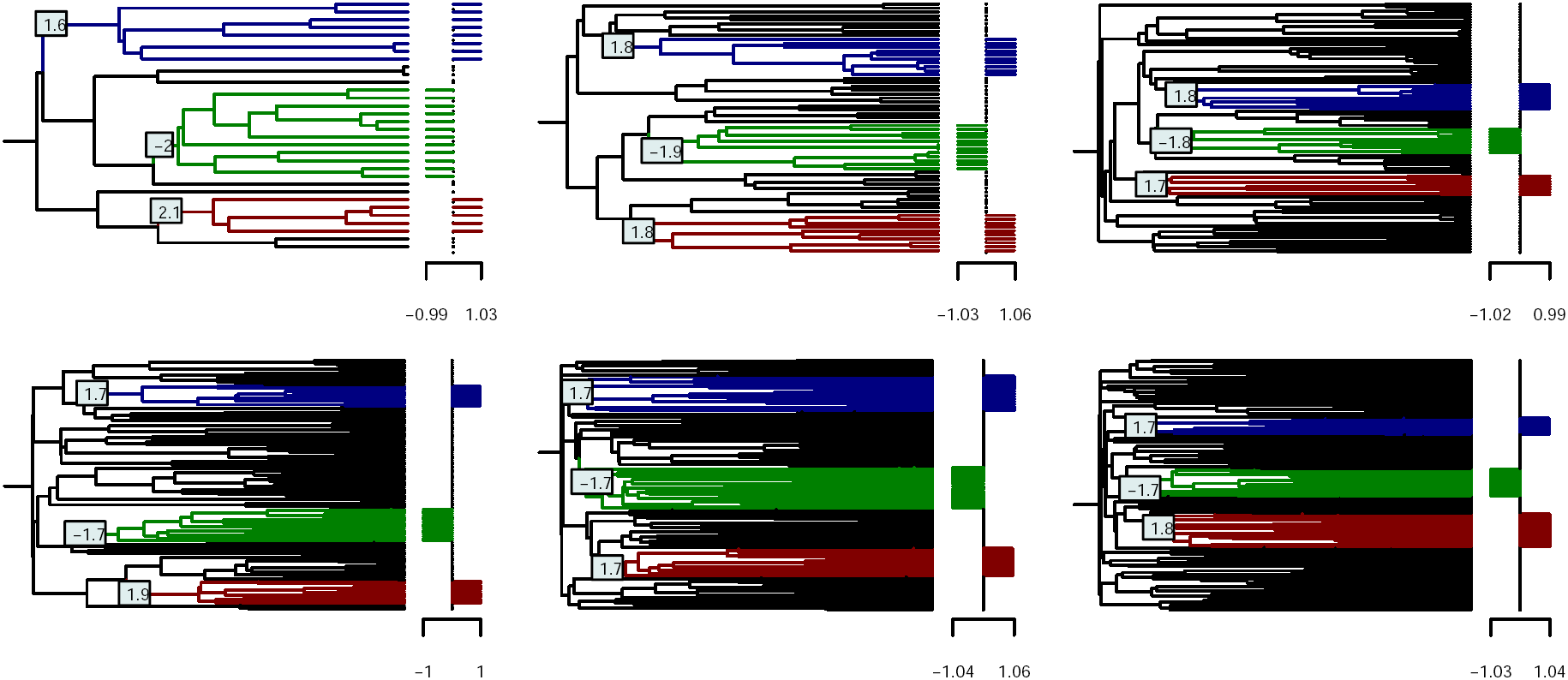
Shifts locations and magnitudes used for the test trees with, respectively, 32, 64, 96, 128, 192, 256 taxa.

### Results

*Number and Amplitude of Shifts*.— We assessed shifts detection accuracy with the Adjusted Rand Index (ARI, Hubert and Arabie 1985) between the true clustering of the tips, and the clustering induced by the inferred shifts (Fig. 5, top). Before adjustment, the Rand index is proportional to the number of pairs of species correctly classified in the same group or correctly classified in different groups. The ARI has maximum value of 1 (for a perfectly inferred clustering) and has expected value of 0, conditional on the inferred number and size of clusters. We use this measure rather than the classical precision/sensitivity graphs as only the clustering can be recovered unambiguously (see Section Model). Note also that when there is no shift (*K* = 0), there is only one true cluster, and the ARI is either 1 if no shift is found, or 0 otherwise (see appendix Note on the ARI).

**Figure 5:**
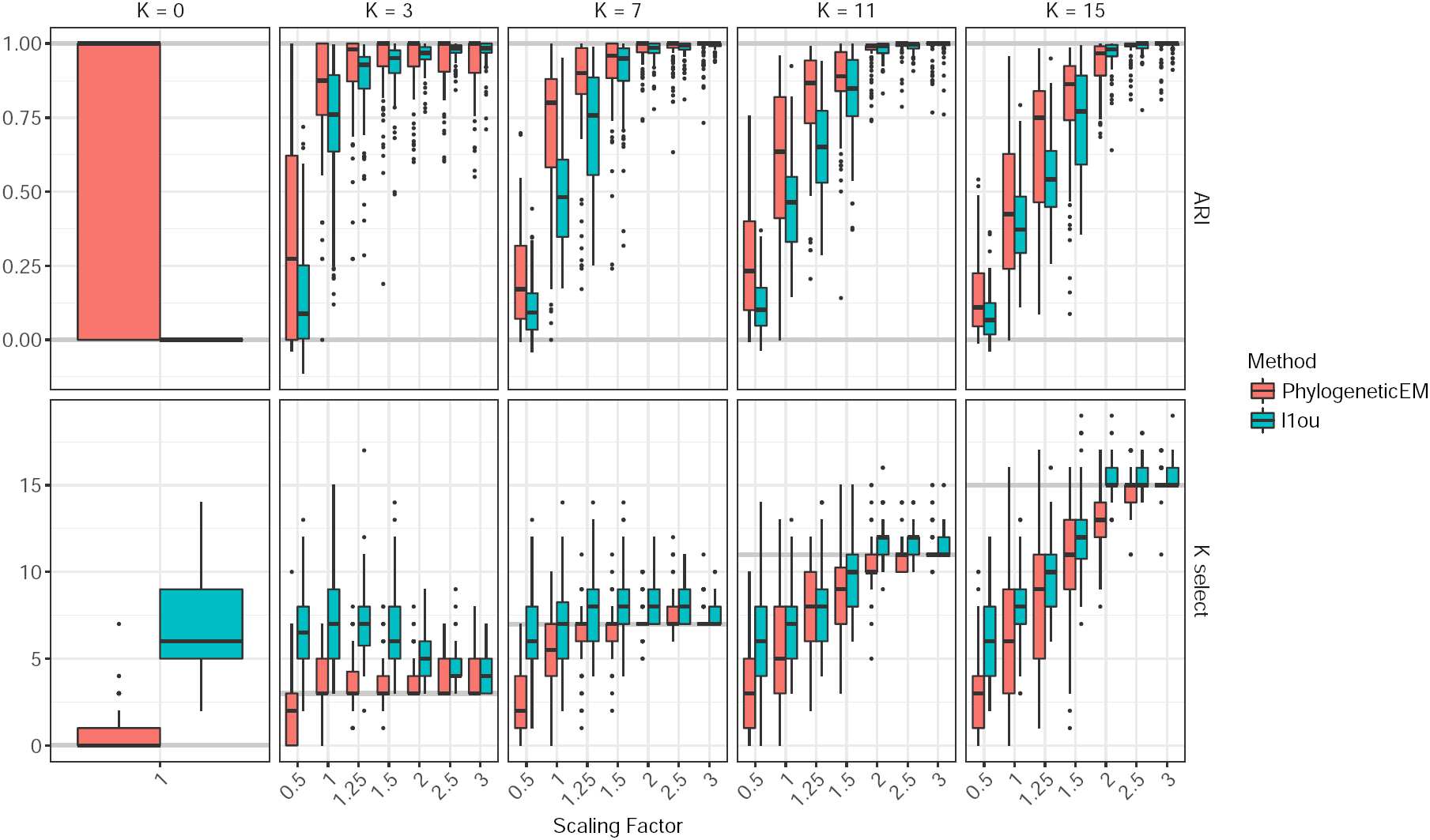
ARI (top) and number of shifts selected (bottom) for the solutions found by **PhylogeneticEM** (red) and *ℓ***1ou** (blue). Each box corresponds to one of the configuration shown in Figure 2, with a scaling factor varying between 0.5 and 3, and a true number of shift between 0 and 15 (solid lines, bottom). For the ARI, the two lines represent the maximum (1) and expected (0, for a random solution) ARI values.

Figure 5 (top panel) shows that, unsurprisingly, both methods detect the number and positions of shifts more accurately when the shifts have higher amplitudes. **PhylogeneticEM** is also consistently better than *ℓ***1ou** when there is a base correlation (here, *r*_*b*_ = 0.4, see section Simulations Studies), which is expected as the independence assumption of *ℓ***1ou** is then violated. The case *K* = 0 (no shift) shows that *ℓ***1ou** systematically finds shifts when there are none, leading to an ARI of 0. More generally, *ℓ***1ou** is prone to over-estimating the number of shifts, even when they have a high magnitude (Fig. 5, bottom) whereas **PhylogeneticEM** is more conservative and underestimates the number of shifts when they are difficult to detect.

*Selection Strength and Model Mis-specifications*.— Our method is relatively robust to model mis-specification (Fig. 6, top). The first panel confirms that, under model A, high values of *α* reduce the stationary variance and lead to higher ARI values and lower RMSEs for continuous parameters (Fig. 6, bottom, leftmost panel). Similarly, scOU (resp. *ℓ***1ou)** achieves high ARI values under well specified models A and B (resp. A and C). The mis-specification of model C (different selection strengths) does not affect scOU much: it has higher ARI dispersion than *ℓ***1ou** but their median ARI are comparable. By contrast, *ℓ***1ou** is severely affected by correlated evolution (model C) and higher levels of correlations lead to significantly lower accuracy, even though group separation is increased (Fig. 3, right). Finally, both methods are negatively affected by correlated selection strengths (Model D), although *ℓ***1ou** seems more robust to this type of mis-specification.

**Figure 6:**
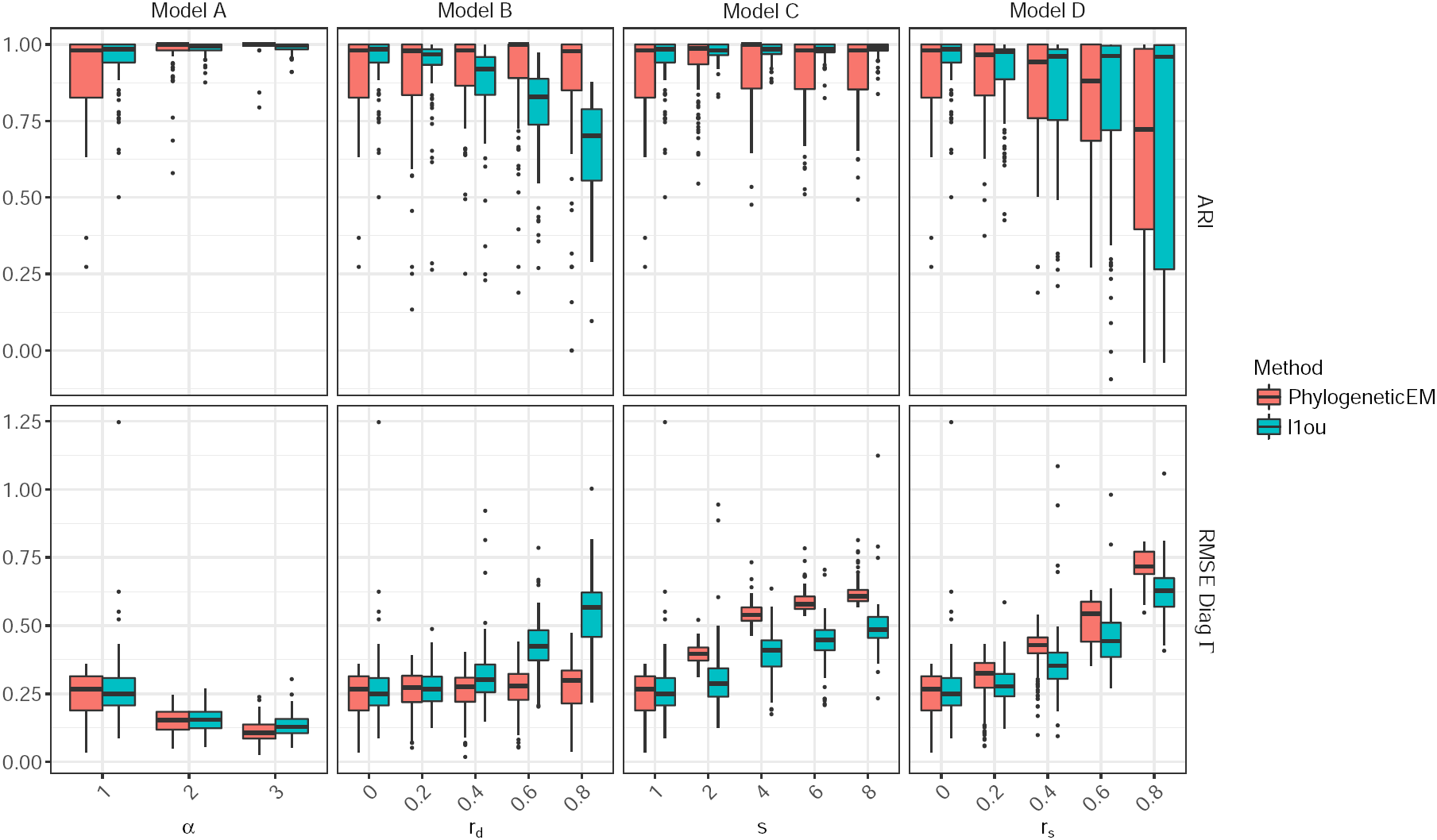
ARI (top) and root mean squared error (RMSE) of the diagonal values of the estimated stationary variance **Γ** (bottom) for the solutions found by **PhylogeneticEM** (red) and *ℓ***1ou** (blue). Each panel corresponds to a different type of mis-specification (except Model A) and the parameters *r*_*d*_, s and *r*_*s*_ control the level of mis-specification, with leftmost values corresponding to no mis-specification. For the ARI, the solid lines represent the maximum (1) and expected (0, for a random solution with the same number and size of clusters) ARI values.

Although shift detection is relatively unaffected by model mis-specification, parameter estimations suffers from it (Fig. 6, bottom, center and right panels). Both *ℓ***1ou** and scOU behave better for model A than for model D and as expected, scOU is not affected by trait correlation (model B) whereas *ℓ***1ou** is. Unequal selection strengths (model C) degrades parameter estimation for both **PhylogeneticEM** and, surprisingly, *ℓ***1ou,** that should in principle remain unaffected. Overall, features of trait evolution not properly accounted for by the inference methods (e.g. correlated selection strengths) are turned into overestimated variances. Note that the quality of the estimation of **Γ** is depends strongly on the estimation of *α*, and could be improved by taking a finer grid for this parameter.

*Number of observations and Computation Time*.— For a given number of shifts, shift detection becomes easier as the number of taxa increases (Fig. 7, left). Furthermore, our method is robust against missing data with detection accuracy only slightly decreased when up to 50% of the observations are missing (Fig. 7, right). Finally, our implementation of the EM algorithm, using only two tree traversals (see appendix EM Inference) and coded in C++, is reasonably fast. Inference takes roughly 15 minutes on a single core on the base 160 taxa tree and less than 45 minutes on the largest simulated trees (256 taxa). *ℓ***1ou** scales less efficiently: it is faster for very small trees (32 taxa) but median running times go up to 20 hours for the large 256-taxon tree. Those long running times were unexpected and higher than the ones reported in Khabbazian et al. (2016). This discrepancy is partly due to the maximum number of shifts allowed, which strongly impacts the running time of *ℓ***1ou**. Khabbazian et al. (2016) capped it at twice the true number of shifts (6 shifts in our base scenario), while we used the default setting, which is half the number of tips (i.e. from 16 to 128 shifts).

**Figure 7:**
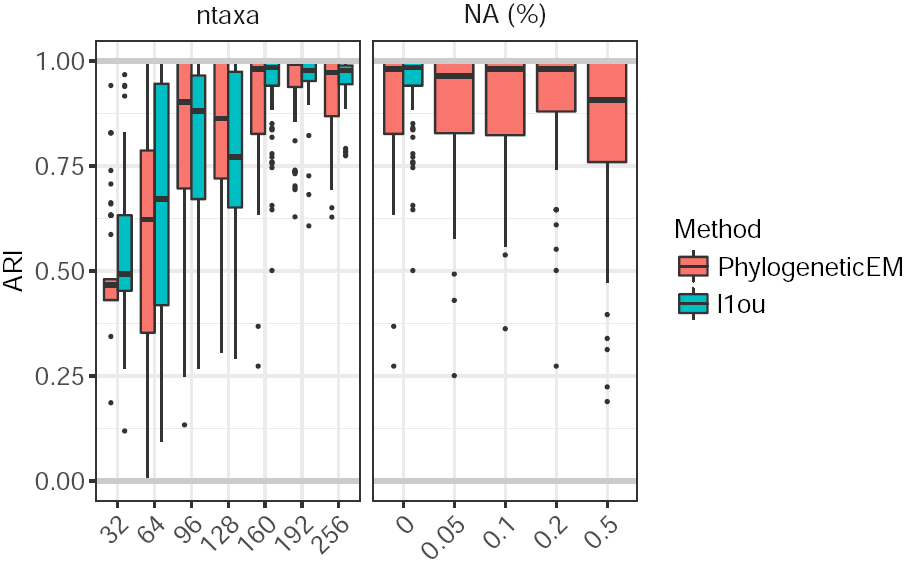
ARI of the solutions found by PhylogeneticEM (red) and *ℓ***1ou** (blue) when the number of taxa (left) or the number of missing values (right) increases. No ARI is available for *ℓ***1ou** when there are missing values as it does not accept them in the version used here, v1.21.

*Impact of pPCA on shift detection accuracy*.— To illustrate how pPCA can both improve and hamper shift detection, we compared **PhylogeneticEM** on raw traits to *ℓ***1ou** on both raw traits and phylogenetic PCs. Figure 9a shows that in our base scenario, with three moderate shifts, pPCA preprocessing slightly decreases performance for low levels of correlations (*r*_*d*_ ≤ 0.2) but drastically improves them for moderate to high correlations levels (*r*_*d*_ ≥ 0.6). Although pre-processing is neutral at moderate correlation levels (*r*_*d*_ = 0.4) with three “easy” shifts, it becomes harmful and degrades the performances of *ℓ***1ou** when the number or magnitude of the shifts increases (Fig. 9b). As expected, **PhylogeneticEM** is unaffected by the pPCA preprocessing, up to numerical issues.

**Figure 8:**
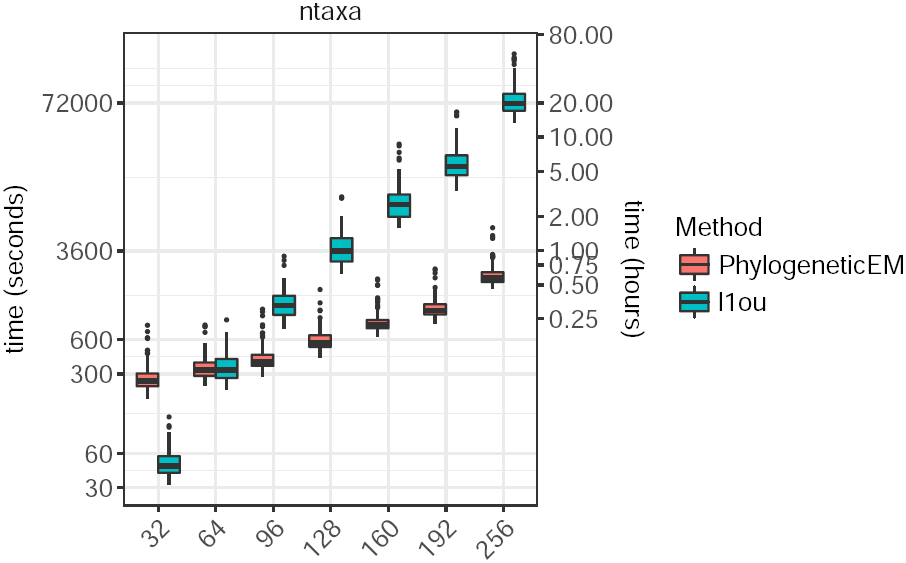
Inference running times (in log-scale) of scOU and *ℓ***1ou**. All tests were run on a high-performance computing facility with CPU speeds ranging from 2.2 to 2.8Ghz.

**Figure 9:**
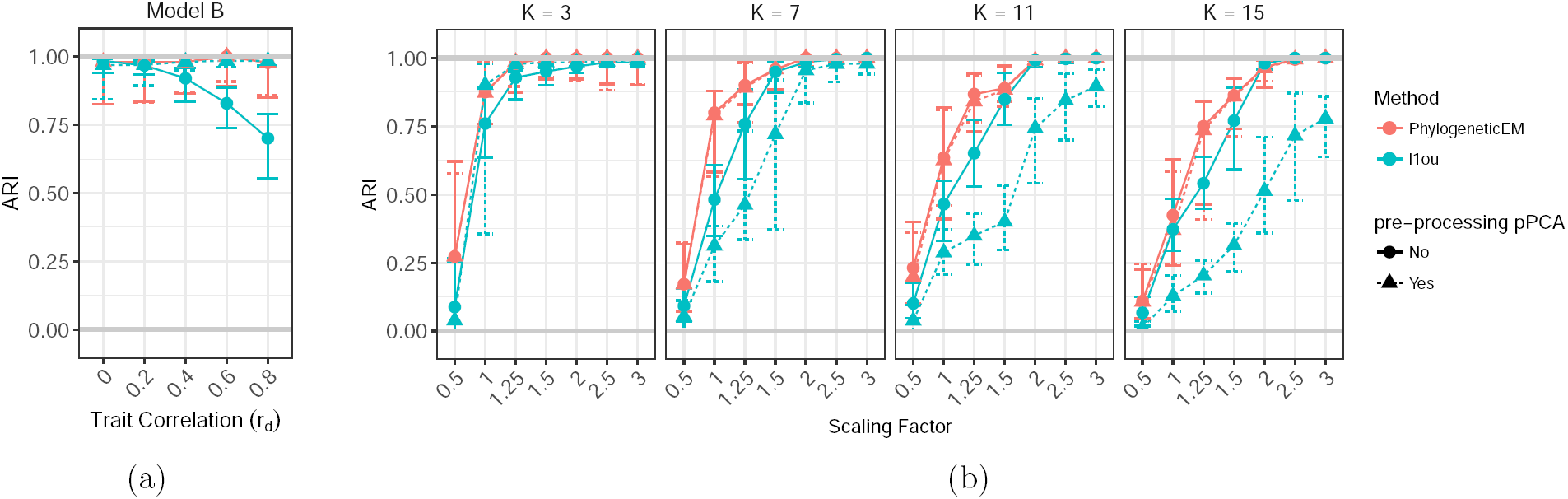
ARI of the solutions found by **PhylogeneticEM** (red) and *ℓ***1ou** (blue), without (solid lines) or with (dotted lines) pPCA preprocessing. (a) Trait correlation (*r*_*d*_) increases from 0 to 0.8. (b) Each box corresponds to one of the configuration shown in Figure 2, and shifts are increasingly large with a scaling factor varying between 0.5 and 3.

## Examples

We used **PhylogeneticEM** to re-analyse two publicly available datasets.

### New World Monkeys

We first considered the evolution of brain shape in New World Monkeys studied by Aristide et al. (2016). The dataset consists of 49 species on a time-calibrated maximum-likelihood tree. The traits under study are the first two principal components (PC1, PC2) resulting from a PCA on 399 landmarks describing brain shape. We ran **PhylogeneticEM** on a grid of 30 values for the *α* parameter. To make this parameter easily interpretable, we report the *phylogenetic half-life t*_1//2_ = ln(2)/*α* (Hansen 1997), expressed in percentage of total tree height. Here, *t*_1/2_ took values between 0.46 % and 277.26 %. We allowed for a maximum of 20 shifts. The inference took 17.56 minutes, parallelized on 5 cores.

The model selection criterion suggests an optimal value of *K* = 4 shifts (Fig. 10, inset graph). The criterion does not show a very sharp minimum, however, and a value of *K* = 5 shifts also seems to be a good candidate. In order to compare our results with that presented in Aristide et al. (2016), we present the solution with 5 shifts (see Fig. 10, left). The solution with 4 shifts is very similar, except that the group with *Aotus* species is absent (in red, see Fig. 10, and supplementary Fig. 14 in Appendix Case Study). Note that, because of this added group, the solution with 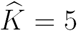has 3 equivalent parsimonious allocations of the shifts (see supplementary Fig. 15 in Appendix Case Study). The groups found by **PhylogeneticEM** (Fig. 10) are in close agreement with the ecological niches defined in Aristide et al. (2016). There are three main differences. First, there is no jump associated with the *Pithecia* species who, although having their own ecological niche, seem to have quite similar brain shapes as closely related species. Second, *Callicebus* and *Aotus* are marked as convergent in Aristide et al. (2016) (in red, right), but form two distinct groups in our model (in pink and red, left). This is due to our assumption of no homoplasy. Finally, the group with *Chiropotes, Ateles* and *Cebus* species (in black) was found as having the “ancestral” trait optimum, while it is marked as “convergent” in Aristide et al. (2016). This is because we did not include any information from the fossil record (not available for brain shape), but instead used a parsimonious solution. Note that the coloring displayed in Aristide et al. (2016) is *not* parsimonious. The two models have the same number of distinct groups.

**Figure 10:**
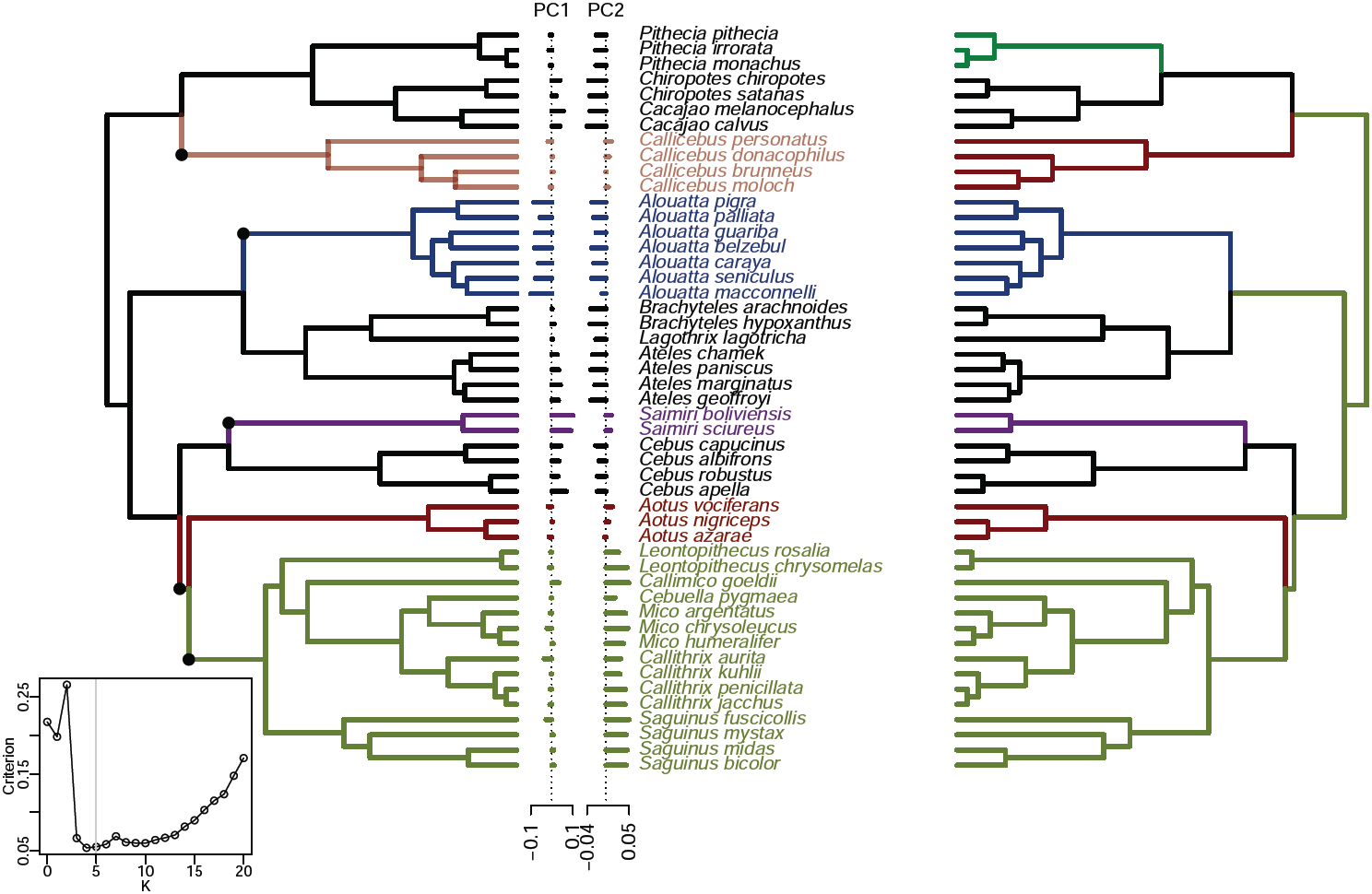
Solution given by **PhylogeneticEM** for *K* = 5 (left) against groups defined in Aristide et al. (2016, Fig. 3) (right), based on ecological criteria including *locomotion* (arboreal quadrupedal walk, clamber and suspensory locomotion or clawed locomotion), *diet* (leaves, fruits, seeds or insects) and *group size* (smaller or larger than 15 individuals). The inset graph shows the model selection criterion. The minimum is for *K* = 4, but *K* = 5 is also a good candidate.

The selected *α* value was found to be reasonably high, with *t*_1/2_ = 12.58%. The estimated correlation between the two PCs was —0.13, confirming that PCA does not result in independent traits.

### Lizards

We then considered the dataset from Mahler et al. (2013), which consists in 100 lizard species on a time-calibrated maximum likelihood tree and 11 morphological traits. We chose this example because of the large number of traits and the high correlation between traits, as all traits are highly correlated (0.82 < *ρ* < 0.97) with snout-to-vent length (SVL).

To deal with the correlation between traits, Mahler et al. (2010, 2013) first performed a phylogenetic regression of all the traits against SVL, retrieved the residuals and then applied a phylogenetic PCA on SVL and the previous residuals, from which they used the first four components (pPC1 to pPC4) for their shift analysis. We first explored how the number of pPCs used can impact the shift detection. Hence we ran **PhylogeneticEM** 11 times, including 1 to 11 pPCs in the input dataset. Each run was done on a grid of 100 values of *α*, with *t*_1/2_ = ln(2)/*α* ∈ [0.99, 693.15] % of tree height, and allowing for a maximum of 20 shifts. It appears that the result is quite sensitive to the number of pPCs included: the selected number of shifts varies from 20, the maximum allowed, to 5 (Fig. 11). When 4 pPCs were used, as in the original study, the estimated covariance matrix **R** contains many high correlations, showing that the pPCs are not phylogenetically independent (Fig. 11).

**Figure 11:**
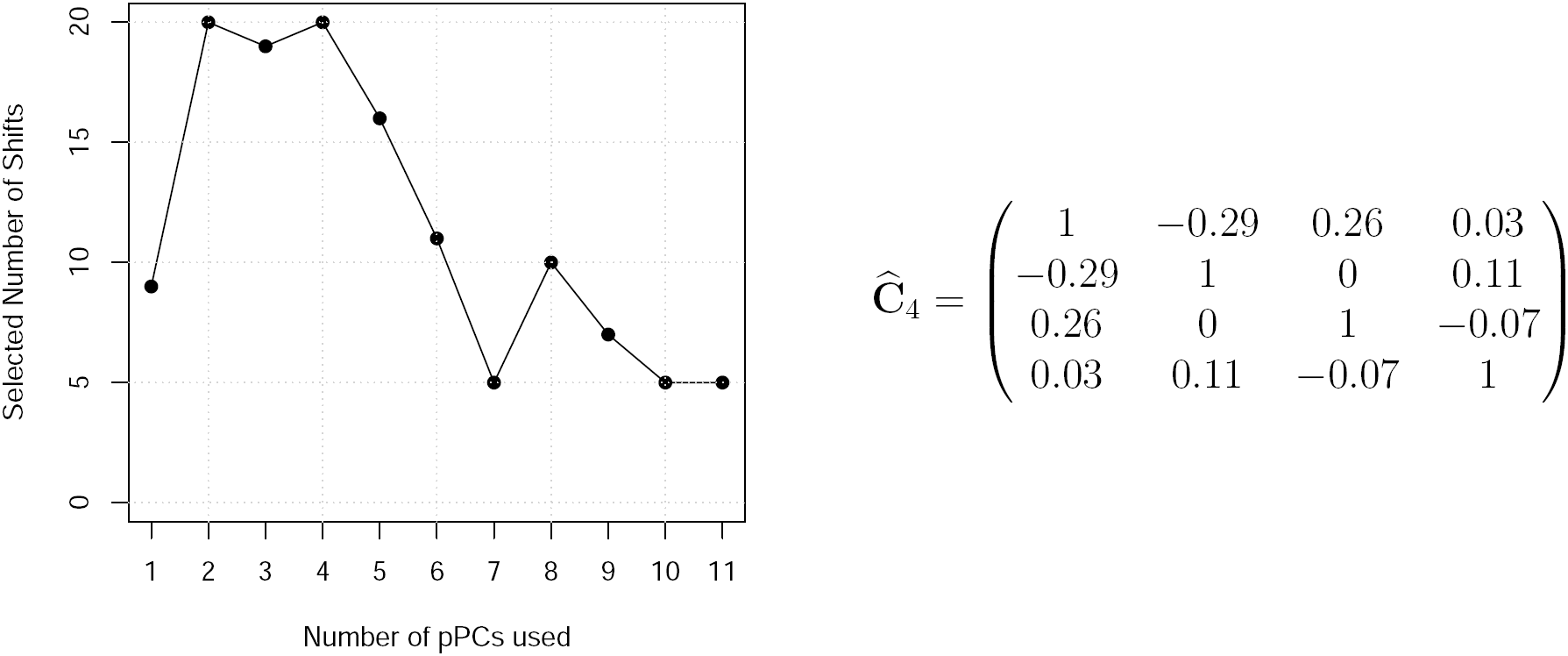
Lizard dataset: selected number of shifts 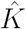given the number of pPCs included in the analysis (left) and estimated correlation matrix between the first four pPCs (right).

To avoid the difficult choice of the number of pPCs, we considered the direct analysis of the raw traits without any pre-processing, and found no shift when running **PhylogeneticEM**. Although the likelihood was found to increase with *K*, the model selection criterion profile was found erratic, suggesting numerical instability. A natural suspect for such instability is the extreme correlation between some traits (0.996 for tibia and metatarsal lengths), which results in bad conditioning of several matrices that must be inverted. To circumvent this problem, we used the two pseudo-orthogonalization strategies described above, running **PhylogeneticEM** on the SVL plus residuals dataset, and on the 11 pPCs, with the same parameters as above. Note that all these transformations use a rotation matrix, so that the likelihood and the least squares of the original or of any of the two transformed datasets are the same. Hence, the objective function, as well as the model selection criterion, should remain unchanged. Still, slight differences were found between the maximized likelihood for each pseudo-orthogonalized datasets. For each value of *K*, we therefore retained the solution with the highest likelihood.

Using the model selection criterion given in Section Statistical Inference, we found 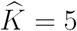shifts, which are displayed in Figure 12, along with the ecomorphs as described in Mahler et al. (2013).

**Figure 12:**
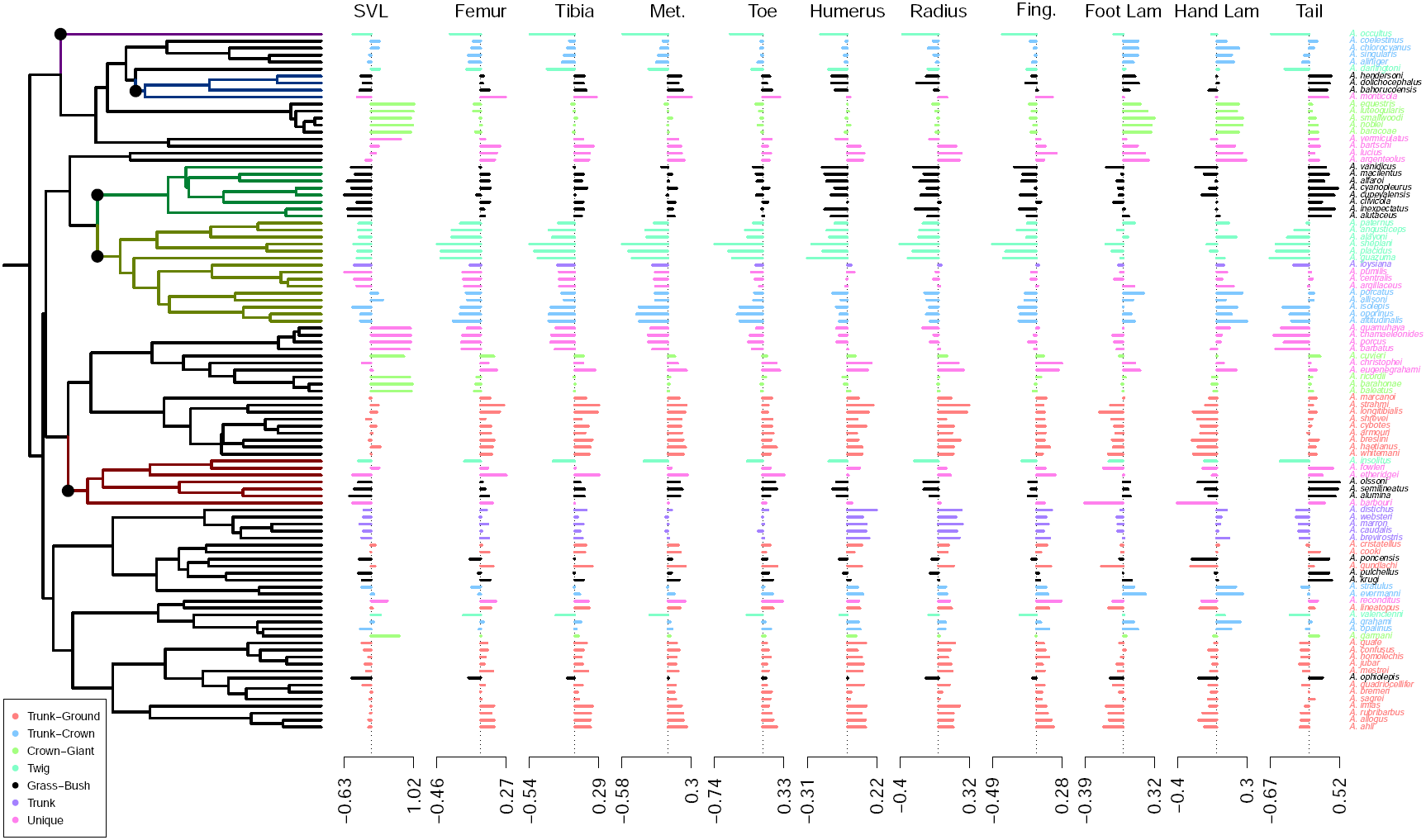
Lizard dataset: solution found by **PhylogeneticEM**. Groups produced by the shifts are colored on the edges of the tree. The species are colored according to ecomorphs defined in Mahler et al. (2013). The traits are the snout-to-vent length (SVL), and the phylogenetic residuals of the regression against SVL of the following traits: femur length, tibia length, metatarsal IV length, toe IV length, humerus length, radius length, finger IV length, lamina number (toe and finger IV), and tail length. The same transformations were used as in Mahler et al. (2010, 2013)

Three of those shifts seem to single out grass-bush *Anolis*, that appear to have a rather small body size, with longer than expected lower limbs and tail, and shorter upper limbs. The two others might be associated with twig *Anolis*, that have smaller than expected limbs and tails. Because of our no-homoplasy assumption, one of those shifts encompasses some species living in other ecomorphs (namely, trunk, trunk-crown and un-classified). The shift, designed to be coherent with the phylogeny, is located on the stem lineage of the smallest clade encompassing the bulk of twig lizards.

### Comments

On both examples (p)PCA does not correct a *priori* for the correlation between the traits in the presence of shifts. In Section pPCA and Shifts we formally proved that it cannot correct for it, actually. As a consequence, any shift detection methods has to account for the correlation between traits.

Still, high correlations between traits may raise strong numerical issues, so PCA can be used as a *pseudo-orthogonalization* of traits, as well as any other linear distance-preserving transformation that would reduce the correlation between them. This does not dispense of considering the correlation between the transformed traits in the model.

The other interest of PCA is to reduce the dimension of the data, which may be desirable when dealing with a large number of traits, such as the original dataset from Aristide et al. (2016). Since PCA does not correct for the right correlation, we have no clue whether or not the dimension reduction performed by PCA is relevant for shift detection, or if it may remove precisely the direction along which the shifts occur. The relevant dimension reduction would consist in approximating the correlation matrix **R** with a matrix of lower rank q < p. This can obviously not be done before the shifts are known, which suggests that shift detection and dimension reduction should be performed simultaneously.

## Discussion

Many phenotypic traits appear to evolve relatively smoothly over time and across many taxa. However, changes in evolutionary pressures (dispersal to new geographic zones, diet change, etc) or key innovations (bipedal locomotion) may cause bursts of rapid trait evolution, coined evolutionary jumps by Simpson (1944). Phenotypic traits typically evolve in a coordinated way (Mahler et al. 2013; Aristide et al. 2015) and a multivariate framework is thus best suited to detect evolutionary jumps. We introduced here an Expectation Maximization algorithm embedded in a maximum-likelihood multivariate framework to infer shifts strength, location and number. Importantly, our method uses Gaussian elimination, just like Fitzjohn (2012), to avoid computing inverses of large variance-covariance matrices and can cope with missing data, an especially important problem in the multivariate setting where some traits are bound to be missing for some taxa. We demonstrated the applicability and accuracy of our method on simulated datasets and by identifying jumps for body size evolution in *Anolis* lizards and brain shapes of New World Monkeys. In both systems, the well-supported jumps occurred on stem lineages of clades that differ in terms of diet, locomotion, group size or foraging strategy (see Aristide et al. 2016 for a detailed discussion) supporting the Simpsonian hypothesis.

### Interpretation Issues

We emphasize that the interpretation of *α* is a matter of discussion. We introduced the scOU in terms of adaptive evolution with a selection strength *α* on the tree. However, the equivalency between OU and BM on a distorted tree suggests that *α* can also be seen as a “phylogenetic signal” parameter, like Pagel’s λ (Pagel 1999). When *α* is small, *ℓ*_*i*_(*α*) ≃ *ℓ*_*i*_ so that branch lengths are unchanged and the phylogenetic variance is preserved. At the other end of the spectrum, when *α* is large, *ℓ*_*i*_(*α*) ≃ 0 for inner branches and the rescaled tree behaves almost like a star tree. However and unlike Pagel’s λ, *α* also dictates how shifts in the optima in the original OU (Δ^*OU*^) are transformed into shifts in the traits values in the rescaled BM (Δ^*ΒΜ*^(*α*)). For small *α*, recall to the optima is weak and shifts on the optima affect the traits values minimally (Δ^*ΒΜ*^(*α*) ≃ 0). By contrast, for large *α*, the recall is strong and shifts on the optima are instantaneously passed on to the traits values (Δ^*ΒΜ*^(*α*) ≃ Δ^*OU*^). Note however that in both cases, the topology is never lost: a shift, no matter how small its amplitude or how short the branch it occurs on, always affects the same species.

Note that if we observed traits values at some ancestral nodes (e.g. from the fossil record), the equivalency between BM and OU would break down: *α* would recover its strict interpretation as selection strength. On non-ultrametric trees, our inference strategy does not benefit from the computational trick to speed up the M step. Similarly to the univariate case, we could write a *generalized* EM algorithm to handle this situation. In Bastide et al. (2016), we used a lasso-based heuristic to raise, if not maximize, the objective function at the M step. It worked quite well, but was much slower. This approach could be extended to the multivariate setting, although with impaired computational burden. Note also that some shifts configuration that are not identifiable in the absence of fossil data become distinguishable with the addition of fossil data. This affects our model selection criterion, which relies on the number of distinct identifiable solutions. Computing this number on a non-ultrametric tree for an OU remains an open problem, and is probaly highly dependent on the topology of the tree.

### Noncausal Correlations

*ℓ***1ou, SURFACE** and **PhylogeneticEM** make many simplifying assumptions to achieve tractable models. Chief among them is the assumption that **A** is diagonal. While *ℓ***1ou** and **SURFACE** both assume independent traits, **PhylogeneticEM** can handle correlated traits through non-diagonal variance matrix **R**. We warn the reader that correlations encoded by **R** are not causal and only capture *coordinated* and non selective traits evolution: i.e. when arm length increases, so does leg length. In order to capture evolution of trait *i in response* to changes in trait *j* (i.e. when arm length strays away from its optimal value, does leg length move away or toward its own optimum) one should rather look at the value of *A*_*ij*_, as was recently pointed out (Reitan et al. 2012; Liow et al. 2015; Manceau et al. 2016). Our simplifying assumptions are justified by various considerations: our focus on inference of shifts rather than proper estimation of **A** and **R,** simulations showing that shift detection is robust to moderate values of off-diagonal terms in **A,** difficulties to simultaneously estimate *α* and shifts even in the univariate case (Butler and King 2004), and computational gain achieved by considering scalar or diagonal **A**. They also suggest that if the focus is on causal correlation in the presence of shifts, a two-step strategy that first detects shifts using a crude but robust model, then includes those shifts in a more complex model, may achieve good performance.

The other simplifying assumption we made is that all traits shift at the same time. It makes formal analysis of identifiability issues and selection of the number of shifts similar to the univariate case, previously studied in Bastide et al. (2016). The assumption is likely to be false in practice, however. Asynchronous shifts are an interesting extension of the model. An ambitious framework would be to build from the ground up a model that allows for different shifts on different traits. It would have to deal with the combinatorial complexity induced by asynchronous shifts, and to use a different selection criterion for the number of shifts. A less ambitious but more pragmatic approach would be a postprocessing of the shifts to select, for each shift, the traits that actually jumped. This would require derivation of confidence intervals for the shift values.

Finally, and unlike **SURFACE** and new version v1.40 of *ℓ***1ou,** our model excludes convergent evolution. This limitation is shared with other shift detection methods such as **bayou** (Uyeda and Harmon 2014) in the univariate case. This exclusion simplifies formal analysis and allows us to borrow from the framework of convex characters on a tree developed in Semple and Steel (2003) but is also likely to be false in practice. A straightforward extension of our method to detect convergence relies again on postprocessing of the shifts: the inferred optimal value of a trait after a shift can be tested to assess whether or not it is different from previously inferred optimal values and warrants a regime of its own.

### Nature of the jumps

We model shifts as instantaneous and immediately following speciation events, like in the punctuated equilibrium theory of Eldredge and Gould (1972). We don’t argue that this is necessary the case. Selection and drift can reasonably be seen as instantaneous over macroevolutionary timescales but by no means over microevolutionary timescales. There is very strong evidence, for example in peppered moths (Cook et al. 2012), that rapid adaptation can happen even in the absence of speciation. However our model does not allow us to distinguish between many small jumps distributed across a branch, one big jump anywhere on that branch and one big jump immediately following speciation, and therefore between punctuated or Simpsonian evolution.

## Acknowledgments

We are grateful to the INRA MIGALE bioinformatics platform **(http://migale.jouy.inra.fr)** for providing the computational resources needed for the experiments.

## PCA: Mathematical Derivations

*Expectation of the estimated Variance-Covariance Matrix*.— Taking 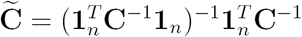, we have that 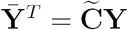, and 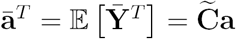. Denote by 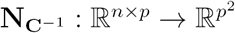the function that to a n *×* p matrix **A** associates the *p × p* matrix **A^*T*^C^−1^A**. We get:

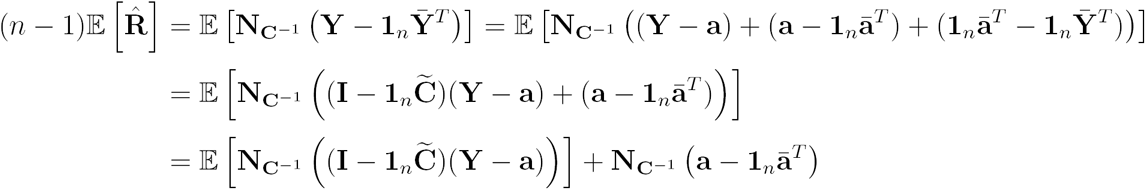

where the two double products cancel out, as 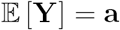. But, for any non-singular symmetric matrix **H,** we have:

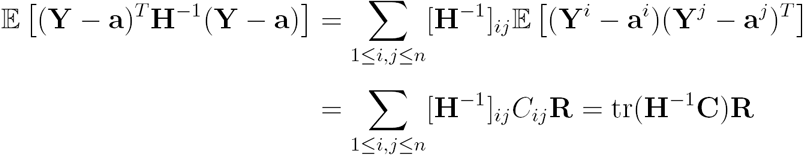

Hence, applying this formula with 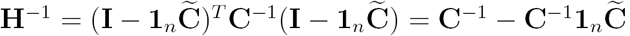, some straightforward matrix algebra manipulations give us:

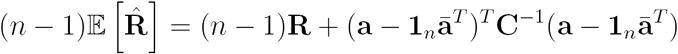

which is the result stated in the text, with 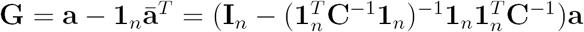.

## PhylogeneticEM package case study: New World Monkeys

In this section, we demonstrate the basic use of the **R** package **PhylogeneticEM** for the analysis of the New World Monkeys dataset (Aristide et al. 2016).

### Loading and Plotting the data

The data have been embedded in the **R** package **PhylogeneticEM,** to be loaded easily. The traits can be plotted on the tree thanks to the function **plot** applied to a void **params_process** object with dimension 2 (Fig. 13).

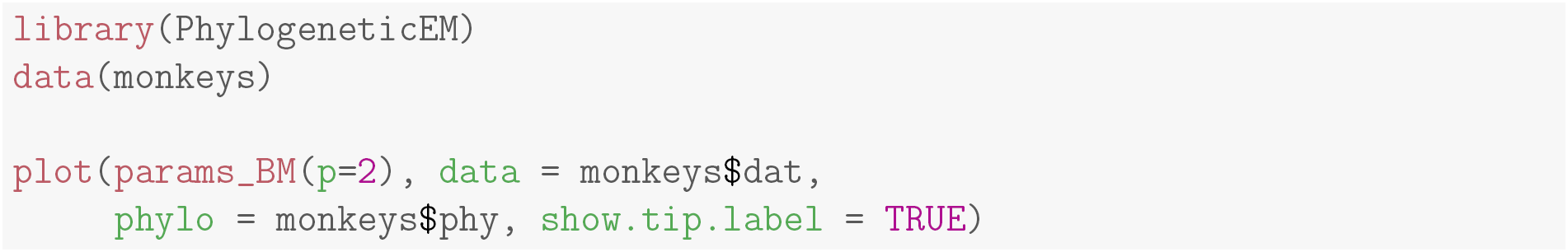

**Figure 13:**
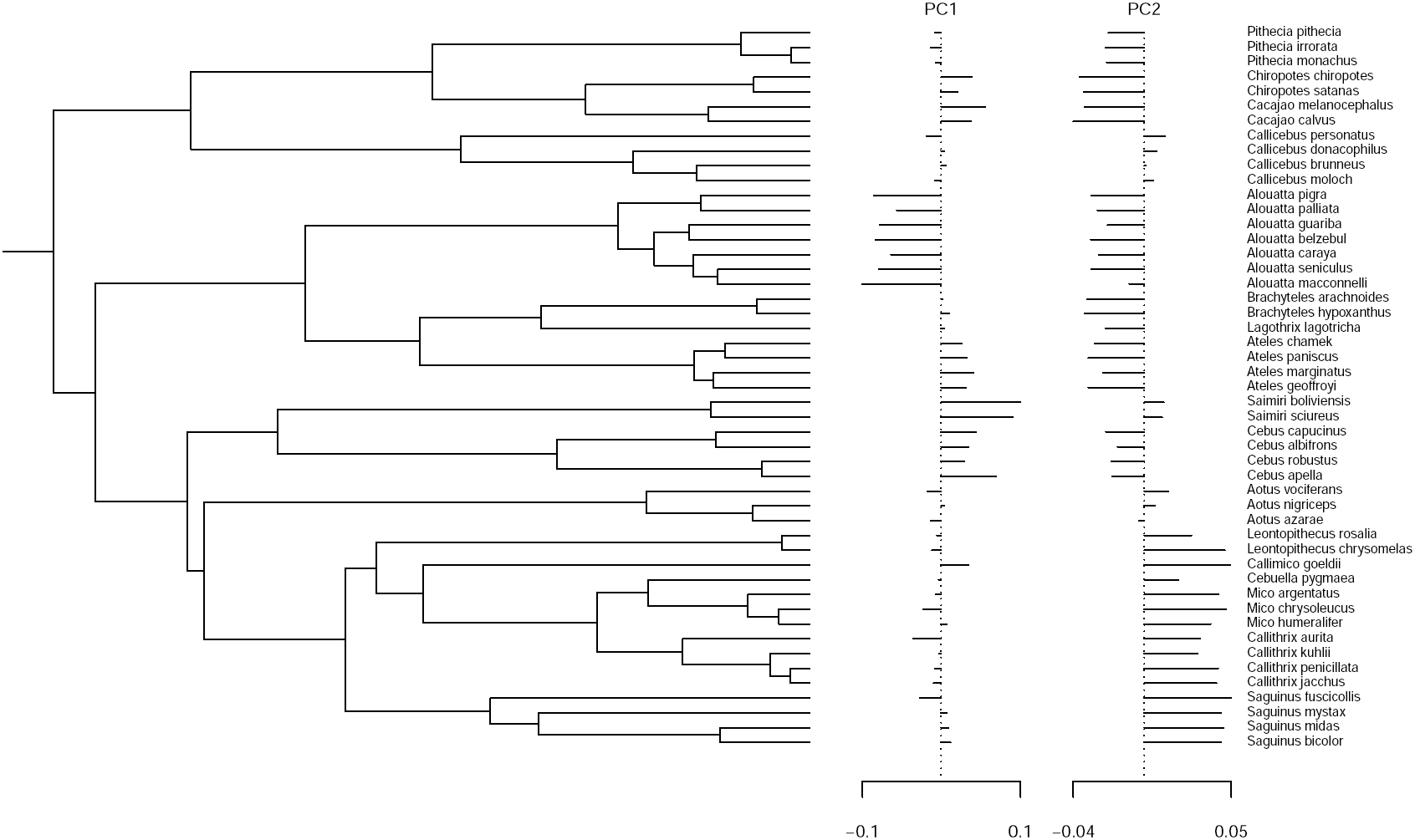
New World Monkey dataset as plotted in **PhylogeneticEM**

This **plot** function inherits from most of the optional arguments of the popular **ape** plot function (here for instance, the optional argument **show.tip.label** is used). Many other graphical parameters can be set by the user, so as to control the output of the function. All the results showed in the main text were produced by the package’s plotting function. The two traits are represented on the right, each with its own scale. Plotting the data on the tree before analyzing it allows us to spot potential errors or outliers.

### Analyzing the data

The automatic shift detection is done using function **PhyloEM**. We show below how the function can be called, using an **scOU** process (with stationary root, the default), for a maximum number of shifts equal to 10, on an automatically chosen grid with 4 values for the selection strength *α*, and parallelized on 2 cores. These parameters were chosen only to demonstrate the function, for this example analysis would run in about one minute. Different parameters were used to obtain the results below and in the main text. There are many more options available to guide the analysis, all described in the manual entry of the function.

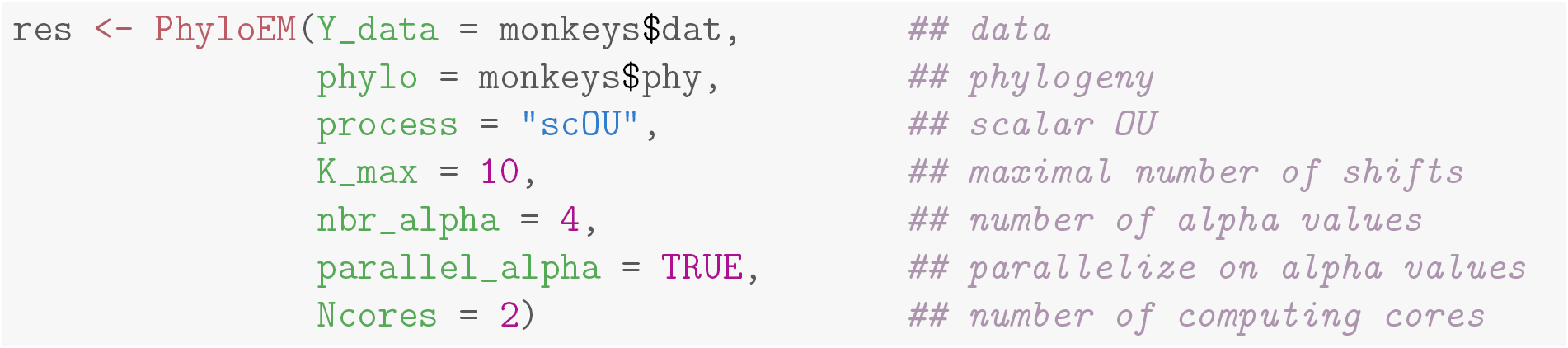

The result is stored in an object of class **PhyloEM,** which has several extractors available (see manual). By default, the plot function draws the maximum likelihood function selected by the method (Fig. 14). The same optional parameters can be used as before to control how the figure should look like.

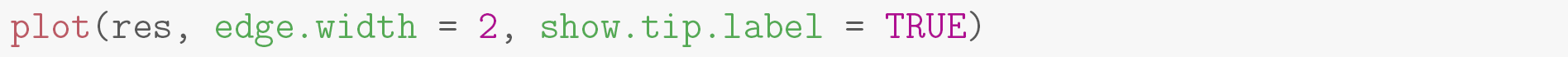

**Figure 14:**
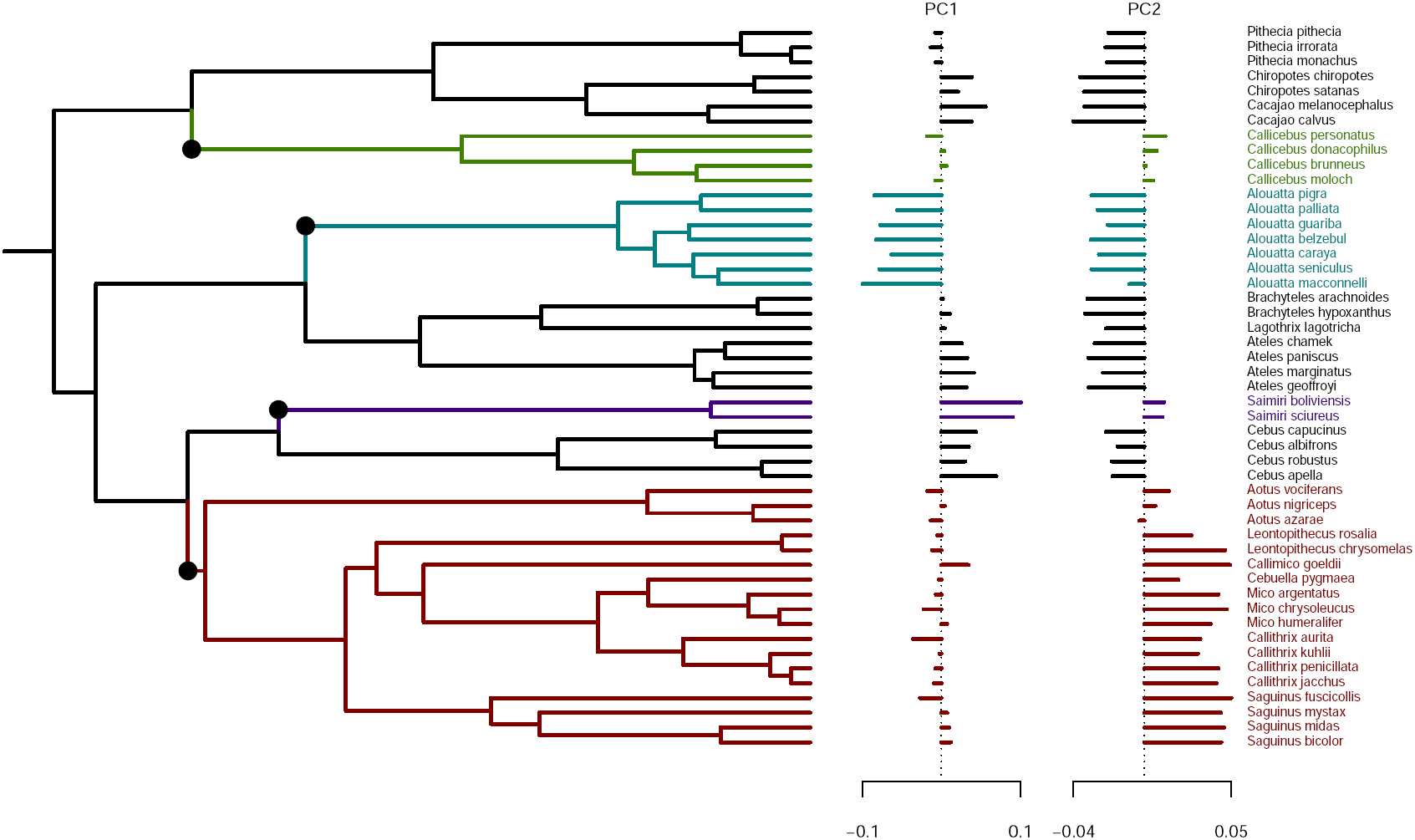
Maximum likelihood solution with 4 shifts selected by the method.

The solution showed in the main text (Fig. 10) has 5 shifts, instead of 4. It can be plotted using the extractor **params_process,** which extracts some inferred parameters from an object of class **PhyloEM**.

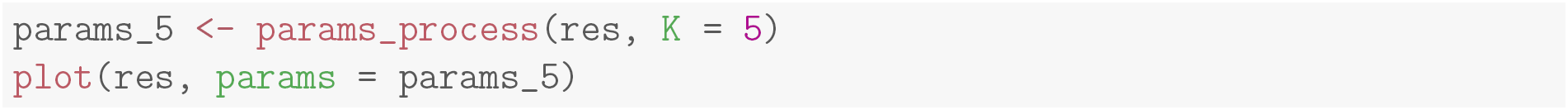

### Plotting Equivalent Solutions

The previous call actually results in a warning being issued: “Warning in **params_process.PhyloEM(res, K = 5):** There are several equivalent solutions for this shift position.” Indeed, as mentioned in the main text, the solution with 5 shifts has three equivalent shift allocations on the branches. These solutions can be found and plotted thanks to the function **equivalent_shifts,** that returns an object that can be visualized (Fig. 15).

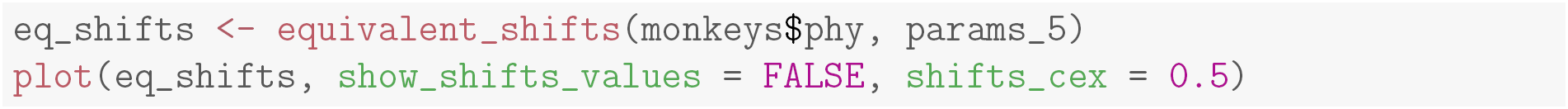

**Figure 15:**
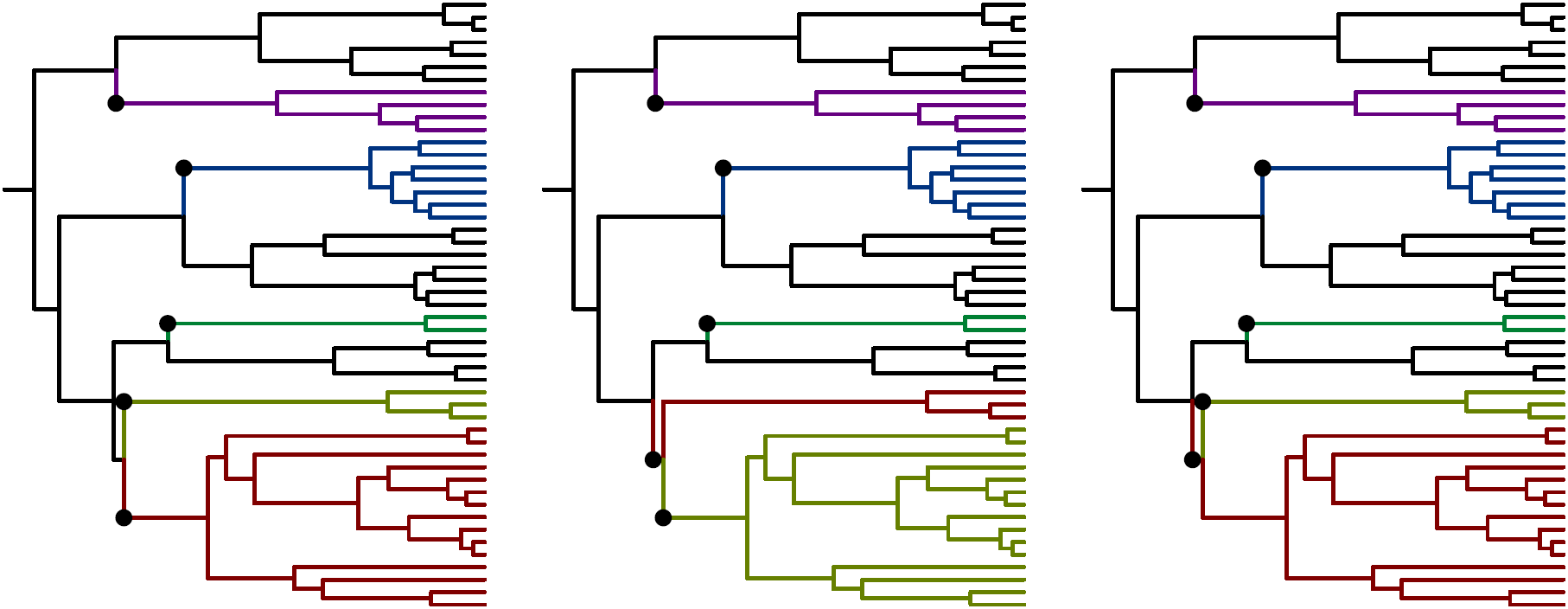
The three equivalent maximum likelihood shift allocations for the solution with 5 shifts.

By default, the shifts values for the first trait is showed for all equivalent solutions. Black is always reserved to the “ancestral state”, and the value λ = *β*_0_ = *μ* of the ancestral optimal value is shown at the root. Here, the three equivalent solutions are quite straightforward, as one configuration has two shifts on sister edges. Note that the clustering of the species at the tips of the tree remains unchanged, while the historic scenario of the adaptive shifts is slightly altered. This ambiguity is inherent to the data. More information to resolve this ambiguity can only come from a prior distribution on shift values, or ideally from fossil data sampled in the right region of the tree.

## EM Inference

This section provides the update formulas for the EM algorithm in Section Statistical Inference. Throughout this section, the superscript *h* refers to the current iteration index, e.g. ***θ*^(h)^** stands for the vector of parameters estimate at iteration h: θ^(*h*)^ = (*µ*^(*h*)^, ∆^(*h*),^ **R**^(*h*)^, **Γ** ^(*h*)^). We denote further by **X** the *N × p* matrix of the traits at all the nodes of the tree, that contains both **Z** and **Y**. In these derivations, nodes are numbered in a preorder, such that the root comes first: *ρ* =1, the internal nodes are numbered from 1 to *m*, and the tips from *m* +1 to *N* = *m* + *n*.

*Conditional expectation of the complete likelihood*.— The EM algorithm mainly deals with 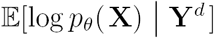, where **Y**^*d*^ is the vector of the observed tips data (that might be missing some values). In our case we have that

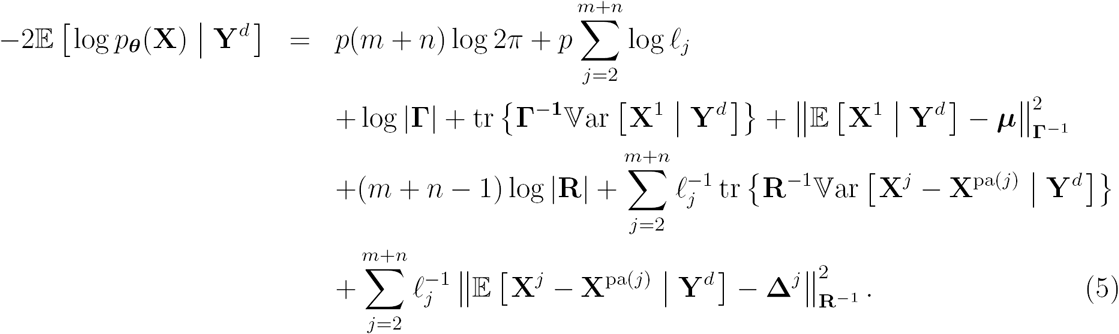

### M step

At the M step, the parameters are updated as the minimizers of (5) evaluated with the conditional moments of the hidden variables given **Y^*d*^**. We get the following updates.

*Root Parameters*.—

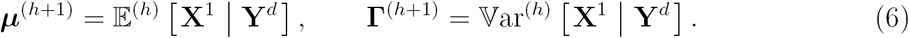

where the conditional moments are obtained as part of the E step, see Equation (8). Notations 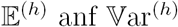denote the moments taken with the law defined by current parameters ***θ*^(*h*)^**.

*Rate Matrix*.—

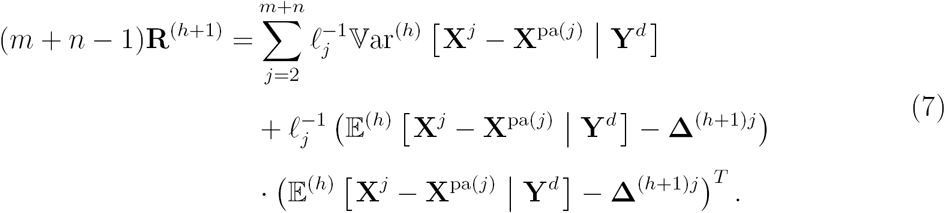

*Optimal Shift Location*.— Only the last term of (5) depends on the shifts so we have to minimize the sum of costs to find **∆**^(*h*+1)^:

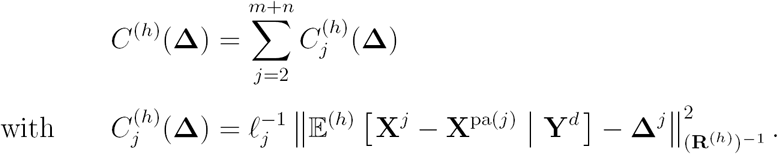

This minimization can be achieved using the same algorithm as in the univariate case (Bastide et al. 2016) to get the optimal shifts allocations and values. Said algorithm essentially sorts the branches in decreasing order of 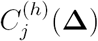 and assigns shifts to the first *K* branches.

### E step

The aim of the E step is to compute the moments of the completed dataset given the observed traits at the tips, namely:

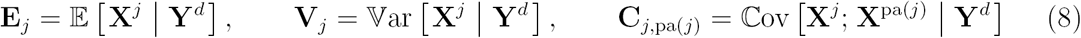

where we dropped the dependency in ***θ***^(*h*)^ for the sake of legibility, but all these moments are indeed taken with the laws given by the current parameters. We do so thanks to an upward-downward recursion on the tree, as described below. This algorithm can apply to a broad classes of Gaussian processes, provided that the moments of the traits at a child node are of the form:

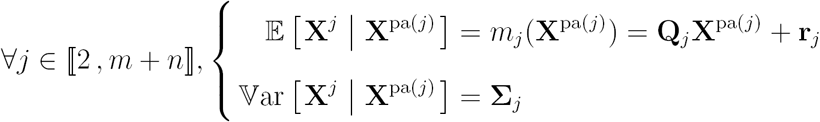

For a BM, we get

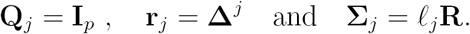

A multivariate OU could also be handled, with:

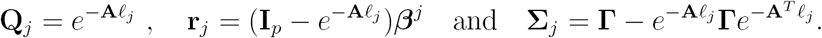

Although we do not use these last formulas here (thanks to the equivalence between OU and BM in our setting), they are implemented in **PhylogeneticEM,** and could be readilly used in an extension of the method to non-ultrametric trees with fossil taxa. To properly handle missing data in a unified framework, we first re-define *ad hoc* inversion and determinant operations that allow us to easily write the degenerated Gaussian likelihood that appears along the way.

*Missing data*.— For a multivariate trait observed at node *i*, define the application 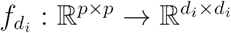that, given a matrix, returns the matrix with only rows and columns corresponding to observed traits. Define also the “pseudo-inverse” 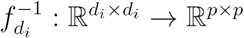that put the observed traits back into their places, and fills the un-defined lines and columns with zeros. This allows us to define a “low-dimensional inverse” as:

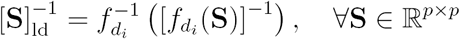

for all **S** such that f_*di*_**(S)** is invertible. We also define a “low dimensional determinant”, as:

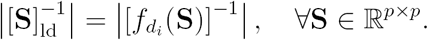

These conventions amount to taking infinite values for the variance-covariance terms of non-observed traits. This allows us to write the following:

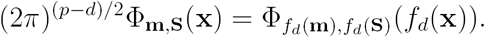

where Φ**_m,S_** denotes the density of a multivariate Gaussian, with expectation vector **m** and variance matrix **S**. That is, we write the density of a *d*-dimensional Gaussian as the density of a *p*-dimensional one, but with the exact same likelihood value, up to a normalizing constant (2*π*)^(*p*−*d*)/2^. If *d* = 0 (no data at one tip), then 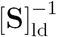 is a matrix of 0, and we take by convention 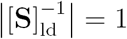, so that 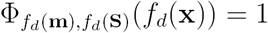.

*Upward recursion*.— For a given node *j* in the tree, we denote by ^*j*^**Y^*d*^** the set of all traits observed at all the tips below node *j*. The aim of the upward recursion is to compute the Gaussian pdf 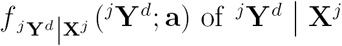, which we write as proportional to a Gaussian density in **a:**

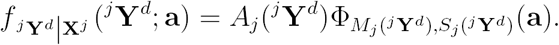

**Initialization:** For each tip i, the observed values **(Y^*d*^)^*i*^** given the vector of values **Y^*i*^** follow a Dirac distribution:

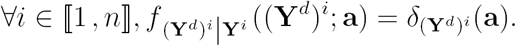

We can express this in the correct format:

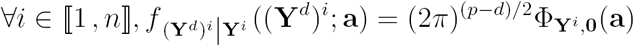

but taking the “low dimensional” inverses and determinants defined above.

**Propagation:** The upward recursion formulas result from the standard properties of the conditional distribution of a multivariate Gaussian distribution plus the fact that *L* daughters of a given node **X^*j*^** are conditionally independent so

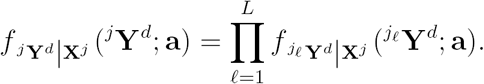

We get

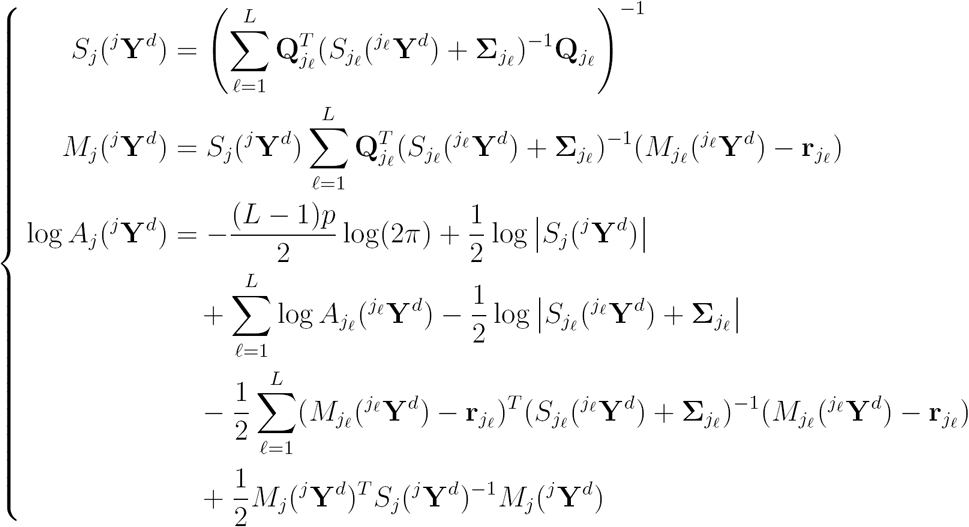

where we keep track of the log of the constant A_*j*_, for numerical accuracy. Remark that we only need to handle the infinite terms properly as described above, using the “low dimensional” inverses and determinants when needed. These terms will disappear as we go up to a node that has at least one tip with some observation for this particular trait. In the pathological case where a trait is never observed, the corresponding term remains infinite throughout the recursion, and hence does not bring any information as to the value of that trait, and does not change the likelihood. The variance of a root non-observed trait is then just the one put a priori in **Γ** (see below).

**Root node and likelihood:** Once at the root, we have 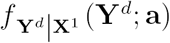, which is the likelihood of the observations given the root state **X^1^** = **a,** and we write:

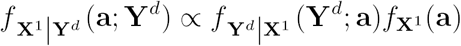

which gives

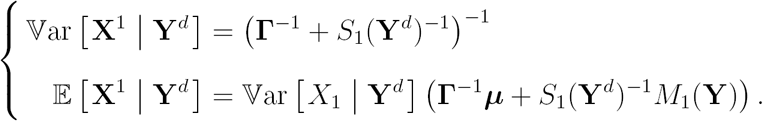

*Downward recursion*.— We now derive a recursion that goes from the root back to the tips to compute the conditional moments required to evaluate (5). Going down the tree, we need to compute, for each node *X*_*j*_, 2 ≤ *j* ≤ *m*, **E**_*j*_, **V**_*j*_ and **C**_*j,pa(j*)_ as in (8). (additionally conditioning on **X^1^** if the root is fixed).

**Initialization:** The initialization of the downward is given by the last step of the upward.

If the root is random, we have

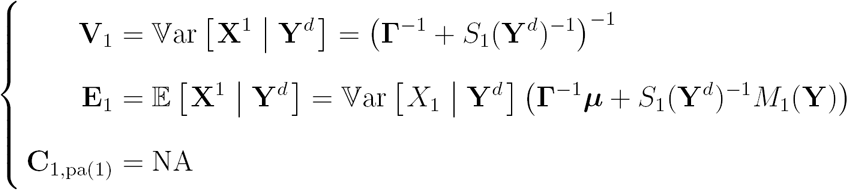

whereas, if we work conditionally to the root, we have 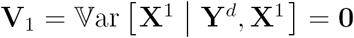, 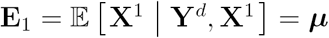and **C_1,pa(1)_** = NA.

**Propagation:** We have

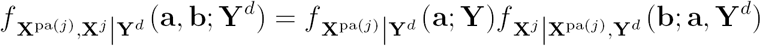

We know the first term from the recurrence, and we can compute the second term thanks to the upward step:

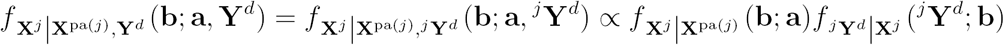

As 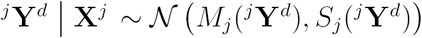and 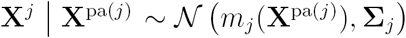, we get

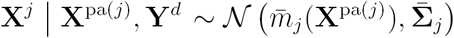

with

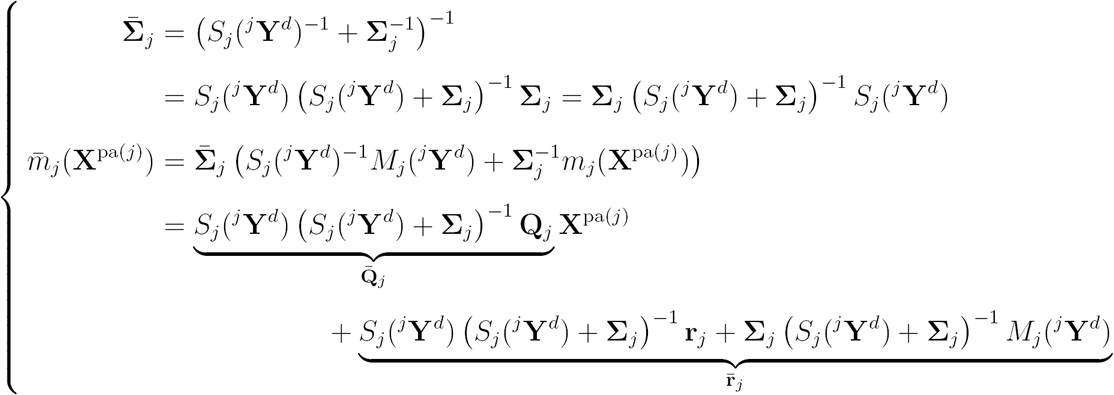

Hence:

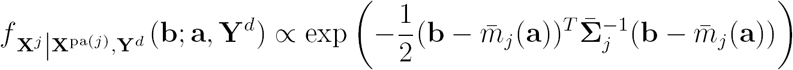

And, as 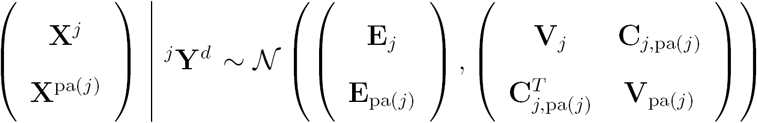, by Gaussian conditioning, we get, for any **a:**

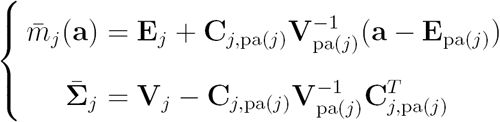

From this we get:

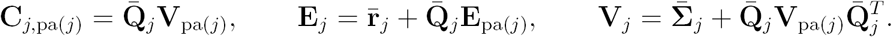

And, finally:

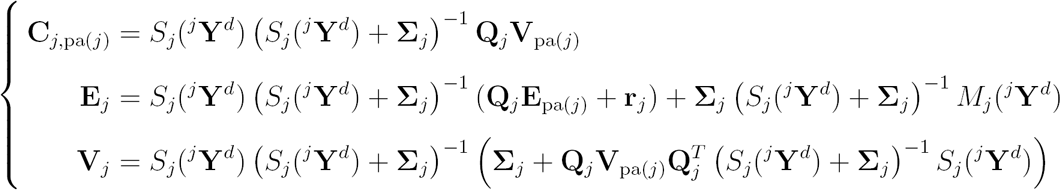

**Missing Data:** In presence of missing data, the downward formulas read

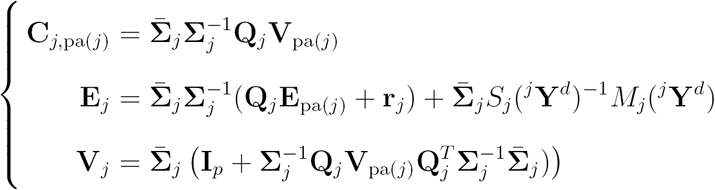

where 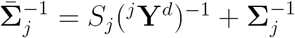can be is computed using the “low dimensional inverse” defined earlier for *S*_*j*_ (^*j*^**Y^*d*^),** if needed.

Remark that theses formulas involve the inversion of two matrices 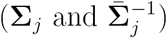, each of dimension *p* (typically small), which is not computationally intensive.

### EM Initialization

Because it is only guaranteed to converge to a local optimum, the EM algorithm is highly sensitive to its starting point. As consequence, it needs to be provided with good initial guesses for the shifts positions and value, as well as the variance matrix **R**. Initial values are determined as follows:

1. Do a lasso regression, assuming all traits are independent, choosing a penalty so that *K* shifts are found.
2. Find the groups of tips created by those shifts, and center each group by its empirical mean.
3. Use the centered data to estimate an empirical variance matrix. This is done using the Minimum Covariance Determinant (MCD) method, with function **covMcd** from package **robustbase** (Rousseeuw et al. 2014).
4. Use this estimated matrix to correct for correlations, before running a lasso again.
5. For this second lasso, choose a penalty that selects for *K* + *K*_lag_ shifts, with *K*_lag_ a fixed value (default to 5). Then, using a Gauss-lasso procedure, select the best *K* shifts (in term of log-likelihood) among those.

This last step can be combinatorially intensive. To keep it fast, we bound the number of trials. It has proven to enhance the results of the algorithm substantially.

### Grid on α

The inference presented above works for the rescaled BM, when the parameter *α* is supposed to be known. In practice, this parameter needs to be estimated. One simple way to do that is to use a grid on *α*. For each value on the grid, one can find an associated estimator, and then find the maximum likelihood estimator of the parameters by taking the best likelihood, for each number of shifts *K*. For instance, we plot below (Fig. 16) the likelihood profile in K for 30 *α* values on a grid, for the New World Monkey dataset (Aristide et al. 2016).

**Figure 16:**
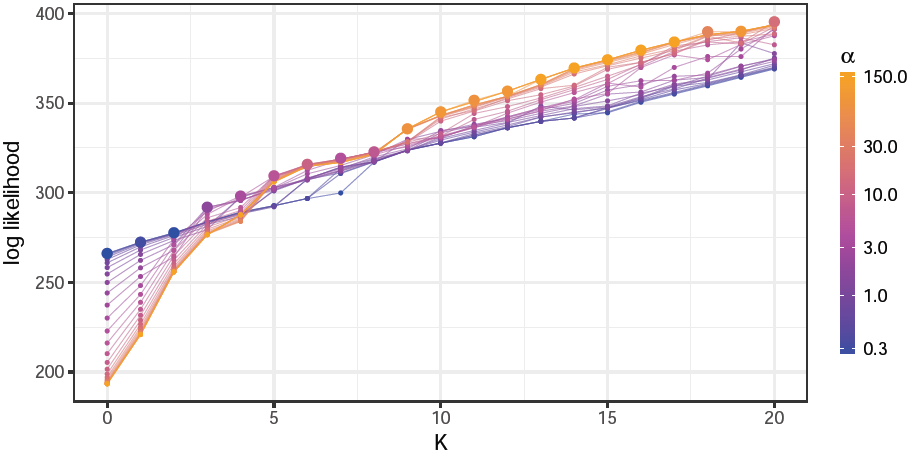
Likelihood profile for all the *α* values, on the New World Monkey dataset. Each colored line represents the likelihood of the solution for a given *α*. The maximum value of the likelihood for each *K* is emphasized. The maximum is not reached by the same value of *α* for each *K*. Colors in log scale.

This grid of *α* values can be provided by the user, depending on some *a priori* knowledge she might have of the problem at hand. If no grid is provided, one is automatically computed, with *n*_*α*_ values, evenly spaced on a log scale ranging between *α*_min_ and *α*_max_. Those extrema values are chosen in the following way.

*α*_min_ The minimum value is chosen so that the maximum phylogenetic half-life (*t*_1/2_ = ln(2)/*α*) is equal to *A* ln(2)*h*, where *h* is the height of the tree, and **A** is a constant, by default equal to 3. This ensures that the lowest *α* makes for a phylogenetic half-life approximately two times as high as the tree. Lower values of *α* would make the process looking too much like a BM.

*α*_max_ The maximum value of *α* is chosen so that the correlations between tips is bounded by *e*^−*B*/2^, with *B* a constant by default equal to 2. This is obtained by noting that the correlation between two tips *i* and *j* for a given trait *k* is given by (for a stationary root):

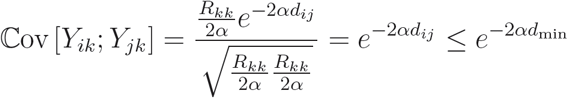

where *d*_min_ is the minimum phylogenetic distance between two tips. Hence we choose *α*_max_ = *B*/(2*d*_min_).

## Simulations Appendices

### Kullback-Leibler Divergences

**Figure 17:**
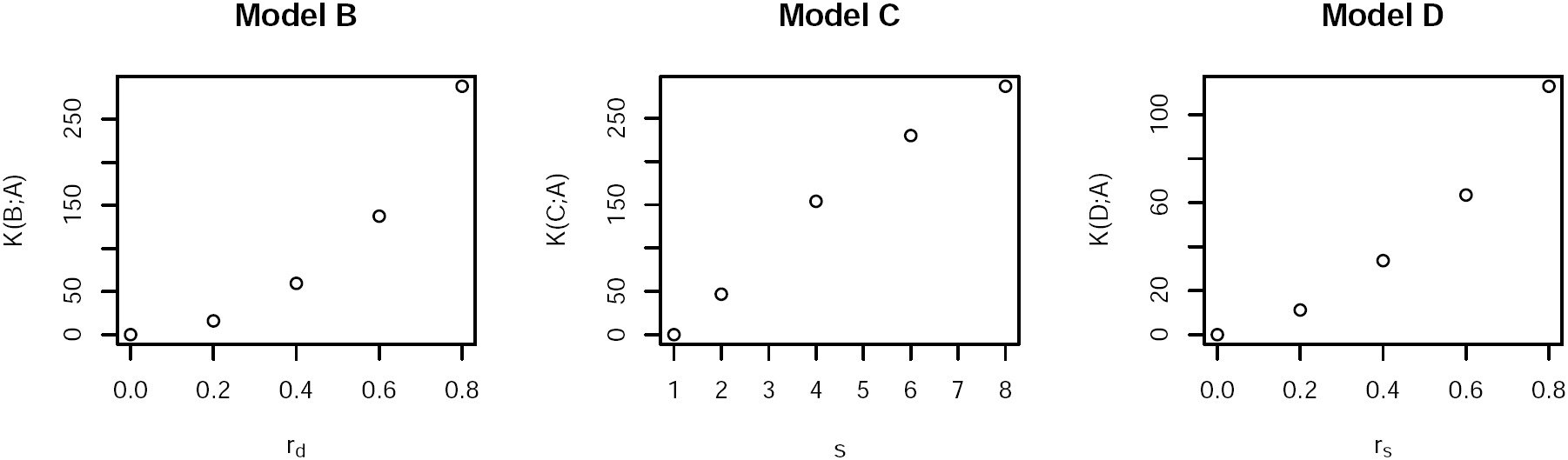
KL divergences from the base model

Denote by **I_*p*_** the identity matrix of size p, **J_*p*_** = **1^*T*^1** the matrix filled with ones, and **S_*p*_** = Diag(*s*^−(*p*+1)/2+*q*^; 1 ≤ *q* ≤ *p*) (so that **|S_*p*_|** = 1). We consider the four following models:

**Model A: A** = **αI**_*p*_ and **R** = σ**^2^I_*p*_**

**Model B: A** = **αI_*p*_** and **R** = **R_*rd*_** = σ**^2^(I_*p*_** + *r***_*d*_(J_*p*_** – **I_*p*_))**

**Model C: A** = **αS**_*p*_ and **R** = σ**^2^S_*p*_**

**Model D: A** = **α**(**I_*p*_** + *r***_*s*_(J_*p*_** – **I_*p*_))** and 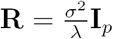

The general formula for a Kullback divergence between two multivariate Gaussian distributions with means *µ*_*i*_ and variances **V_*i*_**(*i* ∈ {1, 2}) is:

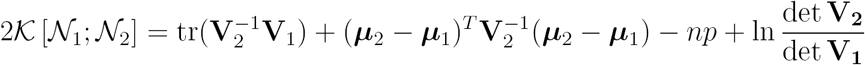

We assume that the root is in the stationary state. From the general formula for a multivariate OU, we derive the form of the variances for these four models (Bartoszek et al. 2012; Clavel et al. 2015):

**General Formula:** 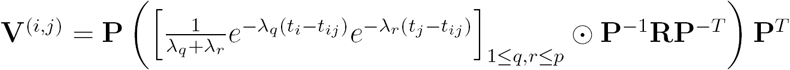, where **P** is the orthogonal matrix of diagonalization of **A,** associated with eigenvalues ^(^λ_l_,…, λ*p*^).^

**Model A:** 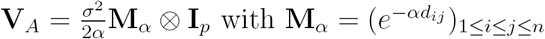

**Model B:** 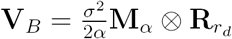

**Model C:** 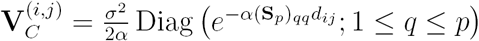

**Model D:** 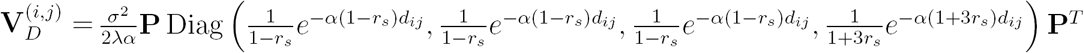

For model C, taking **R** = σ**^2^S_*p*_** ensures that the variances at the tips for all the (independent) traits are equal to 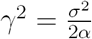.

For model D, the characteristic polynomial of matrix 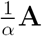 is χ(*Χ*) = (*X* + *r*_*s*_ – 1)^3^(*X* – 3*r*_*s*_ - 1), so we wrote

**A** = *α***Ρ** Diag (1 – *r*_*s*_, 1 – *r*_*s*_, 1 – *r*_*s*_, 1 + 3*r*_*s*_)**P^*T*^**. This leads to a variance at the tips of 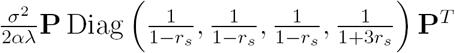. By computing this matrix product (easy linear algebra formula), we find that 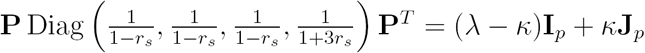, with 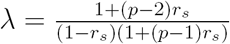and 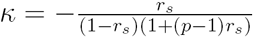. Dividing the variance matrix by a factor *λ* hence ensures that the diagonal variances at the tips are still equal to 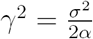.

We can then express the Kullback distance of models B, C and D to model A, using the general formula:

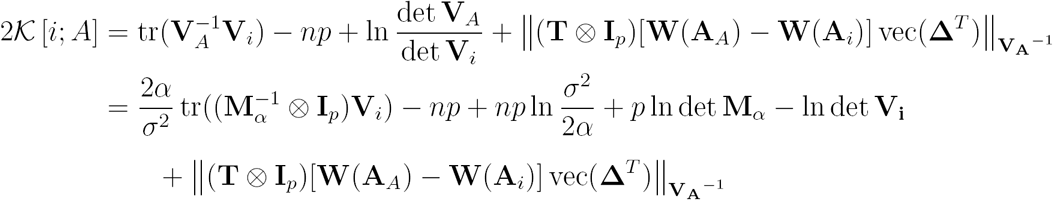

For 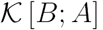, we can get a closed formula that does not depend on the topology (the expectations term cancels out):

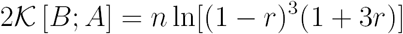

For the two other distances, there are no such nice simplified formula, and the result depends on the topology (even when there are no shifts). To get an idea of the distance when there are no shifts, we computed it on 100 randomly generated trees, and took the mean. With shifts, we computed the distances for the trees and shift positions chosen and shown above.

### Note on the ARI (Hubert and Arabie 1985)

*Partitions*.— Let *S* be a set with n elements, and *U, V* two different partitions of *S*, with respectivelly *R* and *C* groups. Denote by *n*_*ij*_ the number of elements of *S* that are both in groups 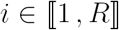of *U* and 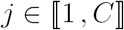of *V*, and by 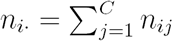(respectively, 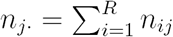) the number of elements of *S* that are in group 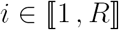of *U* (resp. 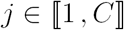of *V*).

*Rand Index*.— We further define:

- *a* the number of pairs of *S* that are in the same groups in both partitions *U* anv *V*,

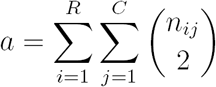
- *b* the number of pairs of *S* that are in different groups in both partitions *U* anv *V*,

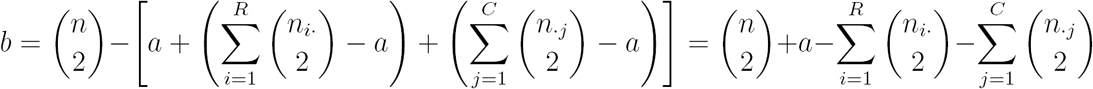

Then the Rand index is defined as the number of agreeing paairs on the total number of pairs:

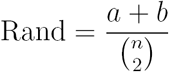

*Adjusted Rand Index*. — Assume that the null model is a generalized hypergeometric models, where the partitions and the number of elements in each group are fixed (i.e. the *n*_*i*_ and *n*_*j*_ are fixed), and the element randomly distributed among them. Then:

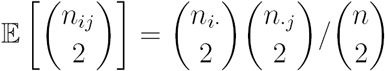

The ARI is then defined as (1 is the maximum value of the Rand index):

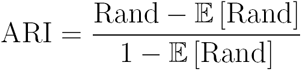

which can be re-written as:

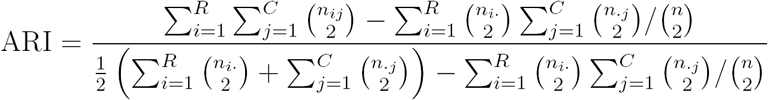

*One class partition*.— Assume that *R* = 1, i.e. that one of the partition has only one class.

Then:

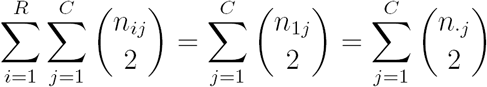

and

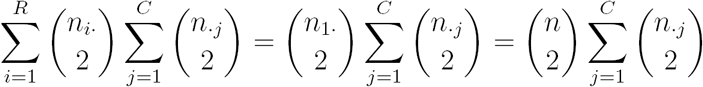

so that ARI = 0. Hence, if one of the true solution or the estimated solution has no shift, then the ARI is automatically equal to 0.

### Supplementary Figures

*Sensitivity / Precision*.— Because only the clustering of the tips induced by the shifts, and not their exact position on the branches of the tree, are identifiable, we used the ARI, rather than sensitivity and recision, to asses methods of shift detection. With this *caveat* in mind, we plot these quantities here for the interested reader. To do that, we removed the 6.53% of solutions that were not identifiable in the results of the methods.

These graphs confirm our conclusions drawn in the main text, with **PhylogeneticEM,** more conservative, having a better precision, along with a similar sensitivity than *ℓ***1ou**. It is interesting to note that, even when the model is violated for **PhylogeneticEM,** the methods keeps a better or similar precision (see e.g. Model C in Fig. 19).

**Figure 18:**
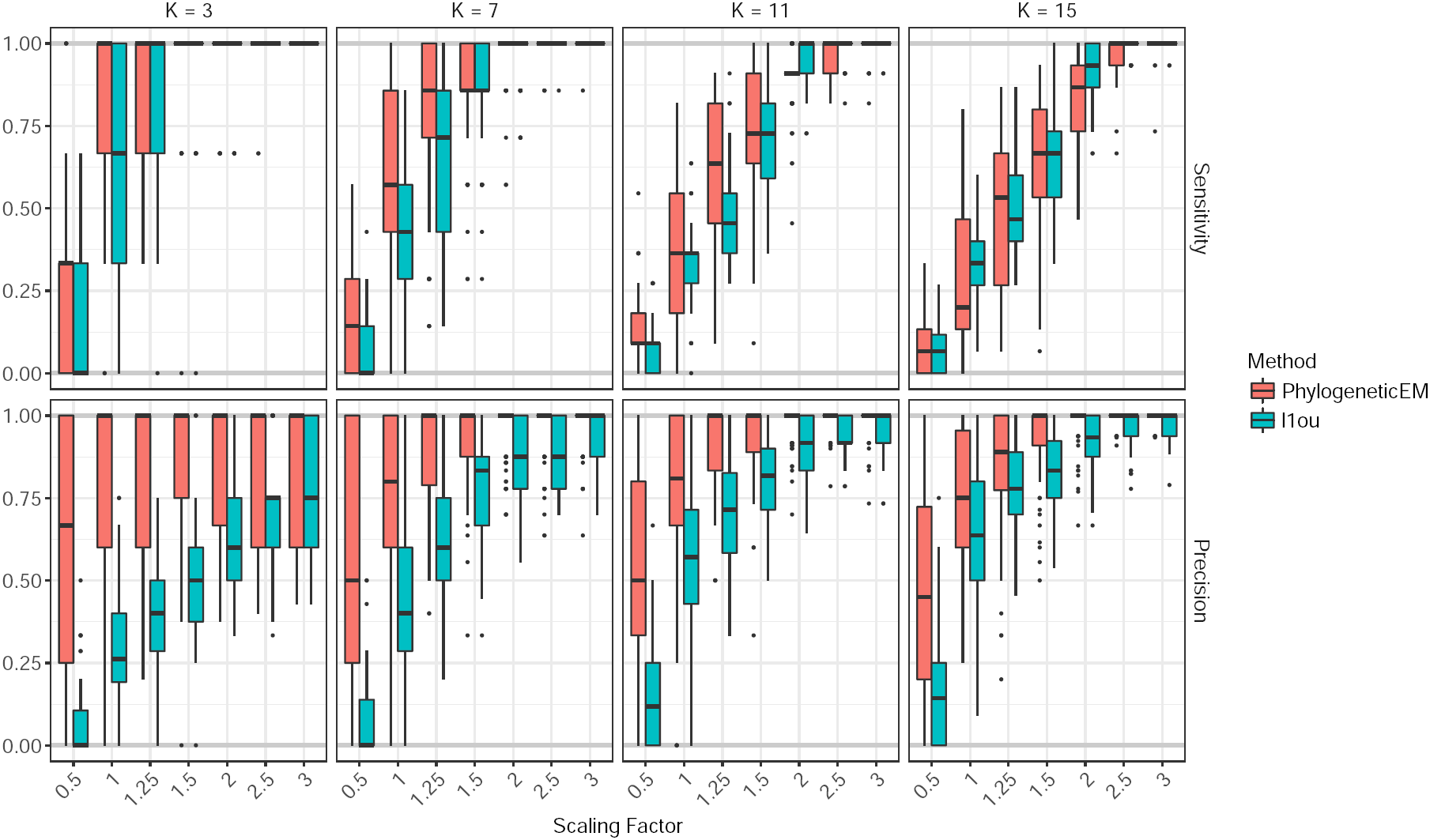
Sensitivity (top) and precision (bottom) for the solutions found by **PhylogeneticEM** (red) and *ℓ***1ou** (blue). Each box corresponds to one of the configuration shown in Figure 2, with a scaling factor varying between 0.5 and 3, and a true number of shift between 3 and 15 (solid lines, bottom).

**Figure 19:**
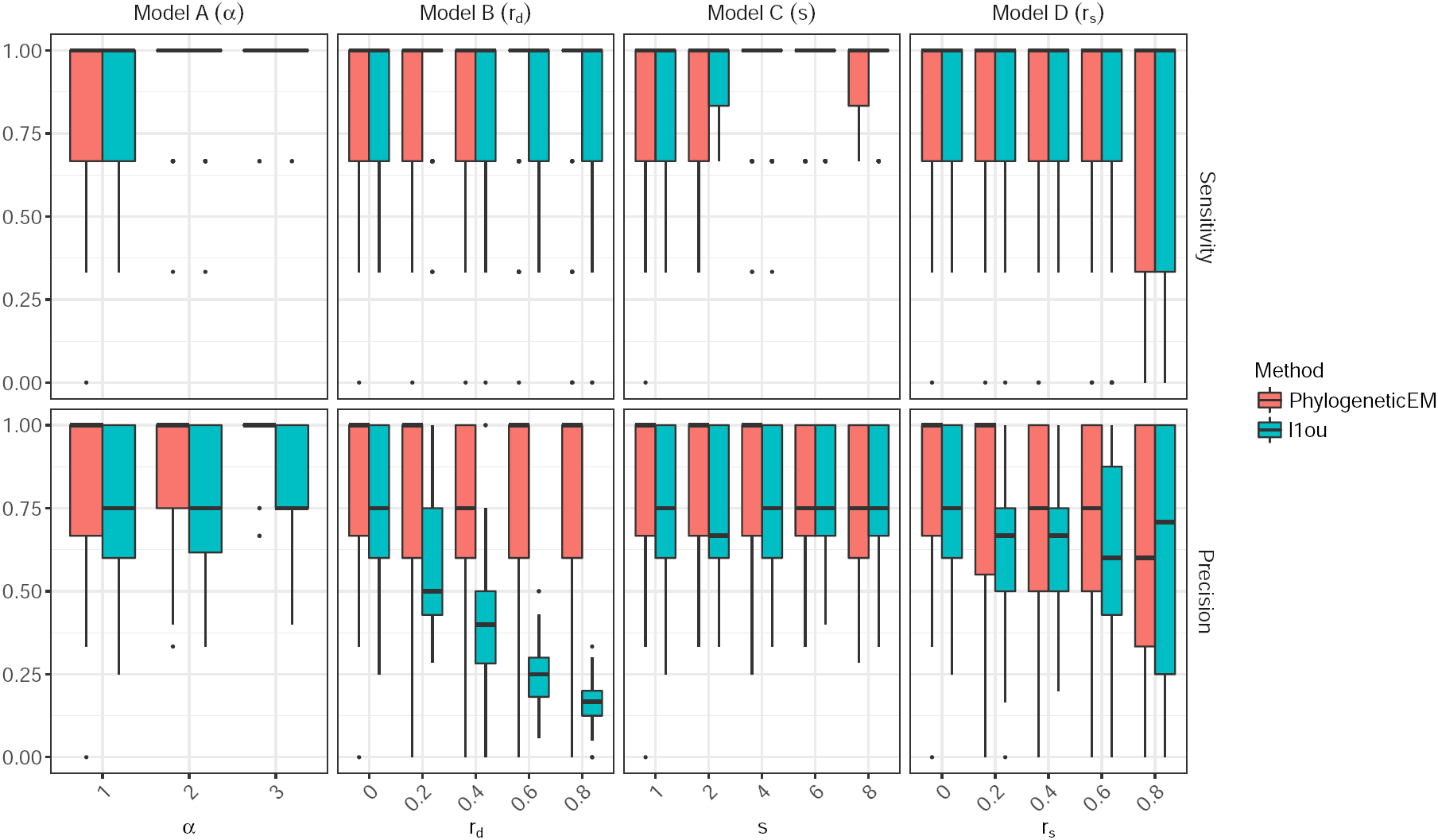
Sensitivity (top) and precision (bottom) for the solutions found by **PhylogeneticEM** (red) and *ℓ***1ou** (blue). Each panel corresponds to a different type of mis-specification (except Model A) and the parameters *r*_*d*_, *s* and *r*_*s*_ control the level of mis-specification, with leftmost values corresponding to no mis-specification. For the ARI, the solid lines represent the maximum (1) and expected (0, for a random solution) ARI values.

